# A novel humanized mouse model to study the function of human cutaneous memory T cells *in vivo* in human skin

**DOI:** 10.1101/490060

**Authors:** Maria M. Klicznik, Ariane Benedetti, Laura M. Gail, Suraj R. Varkhande, Raimund Holly, Martin Laimer, Angelika Stoecklinger, Andreas Sir, Roland Reitsamer, Theresa Neuper, Jutta Horejs-Hoeck, Michael D. Rosenblum, Daniel J. Campbell, Eva M. Murauer, Iris K. Gratz

## Abstract

Human skin contains a population of memory T cells that support tissue homeostasis and provide protective immunity. The study of human memory T cells is often restricted to *in vitro* studies and to human PBMC serving as primary cell source. Because the tisse environment impacts the phenotype and function of memory T cells, it is crucial to study these cells within their tissue. Here we utilized immunodeficient NOD-*scid IL2rγ*^*null*^ (NSG) mice that carried *in vivo-*generated engineered human skin (ES). ES were generated from human keratinocytes and fibroblasts and is initially devoid of skin-resident immune cells. Upon adoptive transfer of human PBMC this reductionist system allowed to study human T cell recruitment from a circulating pool of T cells into non-inflamed human skin *in vivo*. Circulating human memory T cells preferentially infiltrated ES and showed diverse functional profiles of T cells found in fresh human skin. The chemokine and cytokine microenvironment of ES closely resembled that of non-inflamed human skin. Upon entering the ES T cells assumed a resident memory T cell-like phenotype in the absence of infection, and a proportion of these cutaneous T cells can be locally activated upon injection of monocyte derived dendritic cells (moDCs) that presented *Candida albicans*. Interestingly, we found that CD69^+^ memory T cells produced higher levels of effector cytokines in response to *Candida albicans*, compared to CD69^-^ T cells. Overall, this model has broad utility in many areas of human skin immunology research, including the study of immune-mediated skin diseases.

## Introduction

As the body’s outermost barrier, the skin represents a unique and complex immunological organ. As such, healthy human skin contains a large number of CD45RO^+^ memory T cells ^1,2^ that support tissue homeostasis and ensure adequate response to pathogens ^3–5^. A population of resident memory T (T_RM_) cells is found within most tissues where it remains long-term and provides protective immunity after T_RM_ differentiation in response to primary infection ^6,7^. Additionally, T_RM_ may have a protective function in organ transplantation ^8^ and support immuno-surveillance against melanoma ^9^. Cutaneous memory T cells have also been implicated in several diseases, such as cutaneous T cell lymphoma specifically mycosis fungoides ^10,11^.

Generation and maintenance of memory T cells have been extensively studied using murine models ^12,13,13–16^, and significant advances in understanding the role of the skin microenvironment on T cell function and memory development in murine skin have been made ^15,17,18^. Since T cell responses are strongly influenced by the surrounding tissue ^19,20^, and T cells show site-specific functional and metabolic properties ^18,21^, it is crucial to study cutaneous immunity within its physiological compartment *in vivo*. However, direct translation from the murine cutaneous immune system is complicated by fundamental structural differences, as well as a lack of direct correspondence between human and murine immune cell populations ^4,22–24^. Due to technical and ethical limitations, studies of human memory T cell generation have mostly been restricted to *ex vivo* analyses and *in vitro* experiments, and the specific contribution of keratinocyte- and fibroblast-derived signals to cutaneous immunity in human skin remains poorly understood. A better understanding of the requirements of human cutaneous memory T cell recruitment and maintenance in human skin could lead to novel therapies for T cell mediated inflammatory diseases. However, currently existing human immune system (HIS) animal models or skin-xenograft models are complicated by unavoidable inflammation or allo-reactivity. Thus suitable *in vivo* models are required, that faithfully replicate conditions found in human skin under homeostatic conditions to study the requirements for the recruitment and the generation of human cutaneous memory T cell generation and their function *in vivo.*

Skin humanized mice in which immunodeficient mice receive skin grafts from either healthy donors or patients with skin diseases and human peripheral blood mononuclear cells (PBMC) ^25–27^ are currently used to study human inflammatory skin conditions *in vivo*, such as the rejection of skin allografts and xenogeneic graft versus host disease (GvHD) development ^28^. However, T cell recruitment to the skin tissue in absence of inflammation and antigen-specific activation of cutaneous T cells has been much harder to follow. Additionally, if obtained from adult donors skin grafts contain resident immune cells and have a high degree of heterogeneity in terms of immune cell infiltration ^2,25,28,29^, making it difficult to functionally analyze and manipulate discrete skin-tropic T cell populations.

To reduce the heterogeneity found in human skin transplants, bioengineered skin or composite skin grafts were used to study the pathogenesis of inflammatory diseases, such as psoriasis or atopic dermatitis ^30,31^. In these, a sheet of keratinocytes was layered over an *in vitro* generated dermis generated within a fibrinogen or collagen matrix ^32–34^. However, in these models immune cells were applied locally within the engineered skin graft and recruitment of skin-tropic T cells was not studied. Interestingly, data obtained in mouse studies suggested that local skin infection can lead to seeding of the entire cutaneous surface with long lived, highly protective tissue-resident memory T cells, although the highest concentration of these cells occurred at the site of infection ^35^. Repeated re-infections lead to progressive accumulation of highly protective tissue-resident memory cells in non-involved skin ^36^.

Recruitment of human skin-tropic T cells into non-inflamed and inflamed skin is facilitated by several chemokines and cytokines secreted by keratinocytes and fibroblasts ^37–39^. Here we generated a humanized skin mouse model where we utilized mice with human skin engineered only from keratinocytes and fibroblasts to create a reductionist system to study human T cell recruitment to the skin and function within human skin in absence of acute inflammation. Specifically, we used NOD-*scid IL2rγ*^*null*^ (NSG) mice that carried *in vivo-* generated engineered skin (ES) and received human PBMC. This model enabled us to characterize phenotypic changes of circulating memory T cells upon entry of the skin as well as locally restricted antimicrobial responses of human cutaneous memory T cells in absence of infection or inflammation. Additionally, this model offers a new tool to dissect the role of the skin microenvironment in skin immunity *in vivo*.

## Results

### Human T cells specifically infiltrate human engineered skin in a xenograft mouse model

To follow and characterize human T cell recruitment into the human skin *in vivo*, we generated engineered skin (ES) from human keratinocytes and fibroblasts that were isolated from healthy human skin and immortalized ^40^. ES were generated using a grafting chamber as described before ^41^ and allowed to heal and differentiate for a minimum of 30 days (Fig.1a). Consistent with the thorough characterization of the ES by Wang et al. ^41^, Haematoxilin and Eosin (H&E) staining showed that the morphology of the ES was similar to that of normal human skin with an epidermal top layer and an underlying dermal layer (Fig. 1b). The epidermal architecture appeared multilayered and stratified, including a stratum basale, stratum spinosum, stratum granulosum and the stratum corneum, seen as flaking cells in the H&E staining (Fig. 1b, top panel). Human type VII collagen (C7) forms a typical staining-band at the basement membrane zone (BMZ) in immunofluorescence ^42^, which we detected in both primary human skin and the ES. This indicated correct separation of dermis and epidermis of the *in vivo* generated skin (Fig. 1b middle panel). Staining for human cytokeratin 5/6 showed that expression is highest within the stratum basale, in line with the distinct differentiation status of keratinocytes within the epidermal layers of human skin (Fig. 1b, bottom panel).

**Figure 1:**
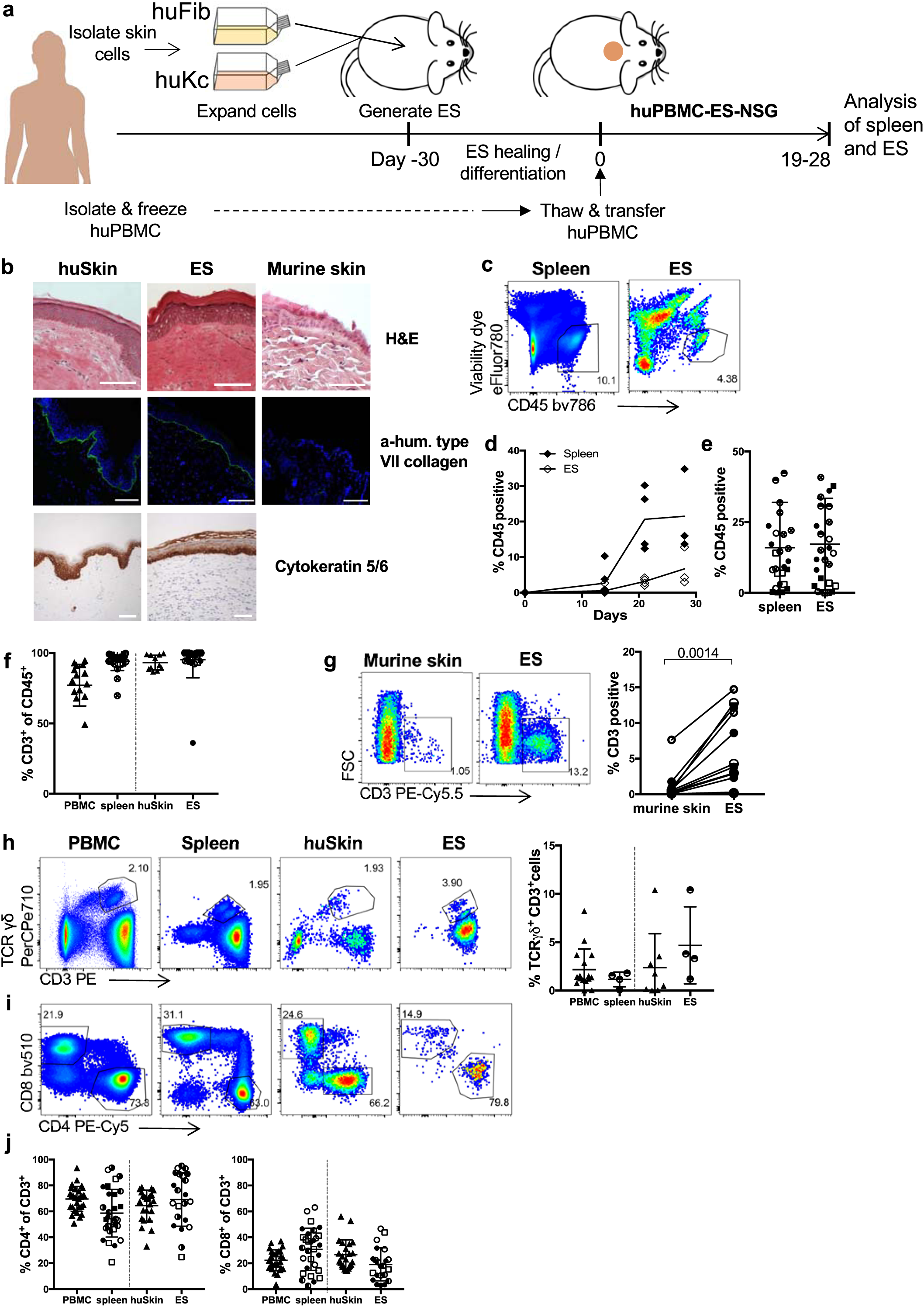
Engineered human skin is preferentially infiltrated by human T cells. (**a**) Schematic of the huPBMC-ES-NSG model. (**b**) ES were generated as in (a) and analyzed by H&E staining and immunofluorescence staining of human type VII collagen on day 72 (upper two panels) as well as immunohistochemical staining of human Cytokeratin 5/6 in ES (lower panel). Murine skin and human skin from a HD served as control. White bar = 100µm (**c-g**) Single cell suspensions of spleen and ES of huPBMC-ES-NSG mice were analyzed by flow cytometry. Each data point represents an individual human donor or experimental mouse. Circles represent data collected from huPBMC-ES-NSG mice using tissue of donor WT85 (female) and squares donor WT70 (male). The different fillings of the symbols indicate independent experiments. (**c**) Representative flow cytometry analysis and (**d**) graphical summary of proportion of human CD45^+^ cell as % of live cells in the lymphocyte gate in paired spleen and ES at indicated time points after adoptive transfer of 2.5×10^6^ PBMC. (**e**) Graphical summary of proportion of CD45^+^ cells of live cells in spleen and ES 18-34 days after PBMC transfer. n=3-6/experiment; cumulative data of 6 independent experiments. (**f**) Graphical summary of the proportion of CD3^+^ cells of live CD45^+^ cells 18-35 days after PBMC transfer; n=3-6/experiment; cumulative data of 6 independent experiments. (**g**) Representative flow cytometry analysis and graphical summary of CD3^+^ percentages in ES and adjacent murine skin 18-35 days after PBMC transfer gated on live lymphocytes. n=3-6/experiment; cumulative data of 3 independent experiments. Significance determined by paired student’s t test; mean ^+^/- SD. (**h**) Representative plots and graphical summary of TCRγδ^+^ and CD3^+^ cells of live CD45^+^ in indicated tissues. (**i**) Representative flow cytometry plots of CD4^+^ and CD8^+^ of CD3^+^CD45^+^ live gated cells (**j**) Graphical summary of CD4 and CD8 expressing cells in human PBMC and skin and spleen and ES, 18-35 days after PBMC transfer gated on live CD3^+^CD45^+^ lymphocytes. n=3-6/experiment; Combined data of 6 independent experiments.

After complete wound healing of the ES, skin-donor-matched PBMC that were isolated and stored in liquid nitrogen until use were adoptively transferred, thus creating a mouse model with a human immune system and ES that we designated huPBMC-ES-NSG (Fig.1a). In previous studies development of xenogeneic GvHD occurred around 5 weeks after adoptive transfer of 10^7^ human PBMC into NSG mice ^43,44^. To delay the development of GvHD we reduced cell numbers to 1.8 – 3×10^6^ /mouse. The weight of experimental mice was monitored throughout the experiments to monitor potential GvHD development. Although we detected no weight loss over a period of up to 87 days following adoptive transfer of 2.5 – 3×10^6^ PBMC (Fig. S1), we limited all experiments to approximately 35 days after PBMC transfer to avoid any potential convoluting effects on our studies.

Following adoptive transfer, we monitored engraftment in the ES and the spleen, which serves as the main peripheral lymphoid organ in NSG mice that lack lymph nodes ^45^. Human CD45^+^ cells were detectable in the spleen after 14 days and in the ES after 21 days (Fig. 1c-d). After a period of 18 - 34 days mean levels of human CD45^+^ cells in spleen and ES were at >18% (Fig. 1e, full gating strategy Fig. S2). The majority of human cells (>94%) in spleen and ES were CD3^+^ T cells (Fig. 1f) and the infiltration of human ES by human CD3^+^ cells was significantly higher compared to adjacent murine skin (Fig. 1g). CD4^+^ and CD8^+^ as well as TCRγδ^+^ T cells engrafted within the spleen and ES at levels comparable to the respective human tissues, PBMC and skin (Fig. 1h-i). The fractions of CD4^+^ and CD8^+^ T cells in spleen and ES reflected the physiological fractions found in human PBMC and skin, respectively (Fig. 1j). This preservation of physiological ratios suggested a specific recruitment process or maintenance mechanism within the ES, similar to human skin. Indeed, T cell-trophic chemokines CCL2 ^46^, CCL5 ^47^, CXCL10 ^48^, CXCL12 ^49^, which support the recruitment of human T cells into human skin ^50^, are secreted within the ES at levels comparable to those of healthy human skin (Fig. 2a).

**Figure 2:**
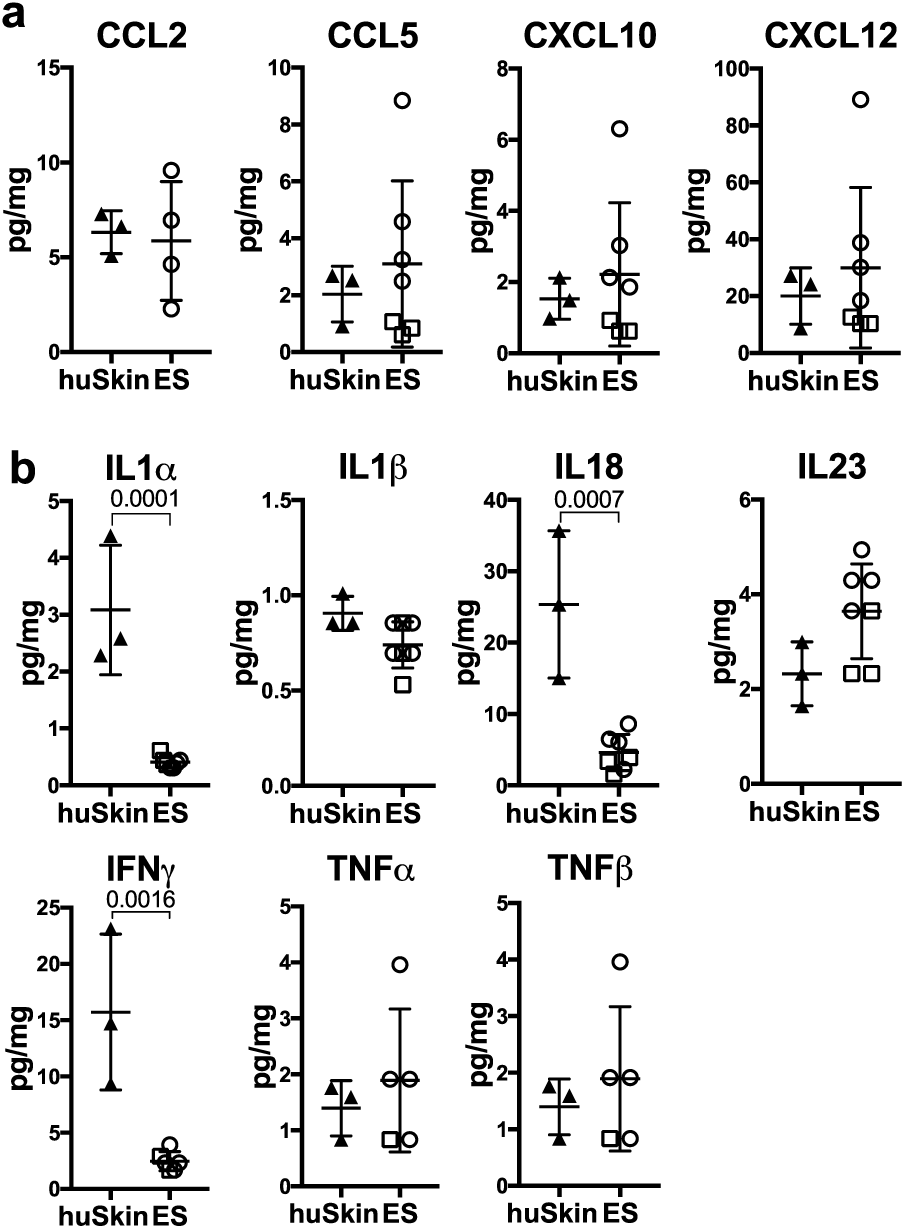
Engineered human skin reproduces chemokine and cytokine levels of non-inflamed human skin. Cytokine and chemokine expression within tissues was determined by bead-based multicomponent analysis of ES from huPBMC-ES-NSG 21 days after PBMC transfer and 3 different healthy human skin donors. Amount of the indicated (**a**) chemokines and (**b**) cytokines per mg skin. Statistical significance determined by student’s t test; mean +/- SD

However, levels of pro-inflammatory cytokines within the ES were equal or even lower to those found in healthy human skin (Fig. 2b), while murine skin lacks these key human chemokines and cytokines. The fact that pro-inflammatory cytokines were not found at increased levels in the ES suggest the absence of an acute inflammation within the engineered tissue. Hence it is unlikely that the preferential infiltration of the human ES over murine skin by human T cells (Fig. 1g) is driven by acute inflammation within the ES, but rather due to the physiological environment within the ES that promotes T cell recruitment and maintenance.

### Engrafted T cells share a skin-homing memory-like phenotype

Since a large proportion of T cells in human skin are memory T cells ^2,25^, we assessed whether this was true for ES-infiltrating CD4^+^ and CD8^+^ T cells (Fig. 3 and Fig. S3). Confirming previous studies of PBMC engraftment in NSG mice, we found that human CD4^+^ as well as CD8^+^ T cells isolated from spleens of huPBMC mice did not express markers of naïve T cells such as CCR7 and CD45RA despite being present in the ingoing PBMC population ^43^, and the vast majority of T cells within the ES had also assumed a CCR7^-^ and CD45RA^-^ memory phenotype (Fig. 3a and Fig. S3a). Similar to the transferred PBMC, the spleen contained CD4^+^ and CD8^+^ T cells that expressed cutaneous leukocyte antigen (CLA), a glycan moiety that promotes skin-homing ^2^. Consistent with this, CLA^+^ T cells accumulate within human skin ^2^ and these cells were also significantly enriched in the ES compared to spleen (Fig. 3b and Fig. S3b). This indicated preferential recruitment or maintenance of skin-tropic memory T cells within the ES. In line with this, IL7 and IL15, two cytokines that support memory T cell function and maintenance in human skin ^51–53^, were found at equal levels within the ES and healthy human skin (Fig. 3c). Upon entering the ES, both CD4^+^ and CD8^+^ T cells upregulated CD69 expression (Fig. 3d,e and Fig. S3c,d), a marker closely associated with tissue residency of human skin T cells ^25^. Interestingly, this was more pronounced within the CD4^+^ compared to the CD8^+^ T cell population. Consistent with CD69 expression, these skin-homing CLA^+^ T cells expressed CCR6 a chemokine receptor characteristic for tissue-resident memory cells ^54^ (Fig. 3d,f and Fig. S3c,e). Additionally, a small fraction of cutaneous T cells also expressed CD103 (Fig. 3g,h and Fig. S3f,g), another marker of human skin T_RM_ ^25^. It remains to be determined whether these T_RM_-like cells are truly resident and maintained long-term or transiently upregulated markers of tissue residency. By contrast, circulating CD62L^+^ memory T cells could be identified in both spleen and ES among both T cell subsets, representing central memory T cells. (Fig. 3i, j and Fig. S3h, i). Taken together these data indicate that upon entry into the ES, circulating CD4^+^ and CD8^+^ T cells reconstitute a skin-tropic T cell population and assume a memory-like phenotype within the ES.

**Figure 3:**
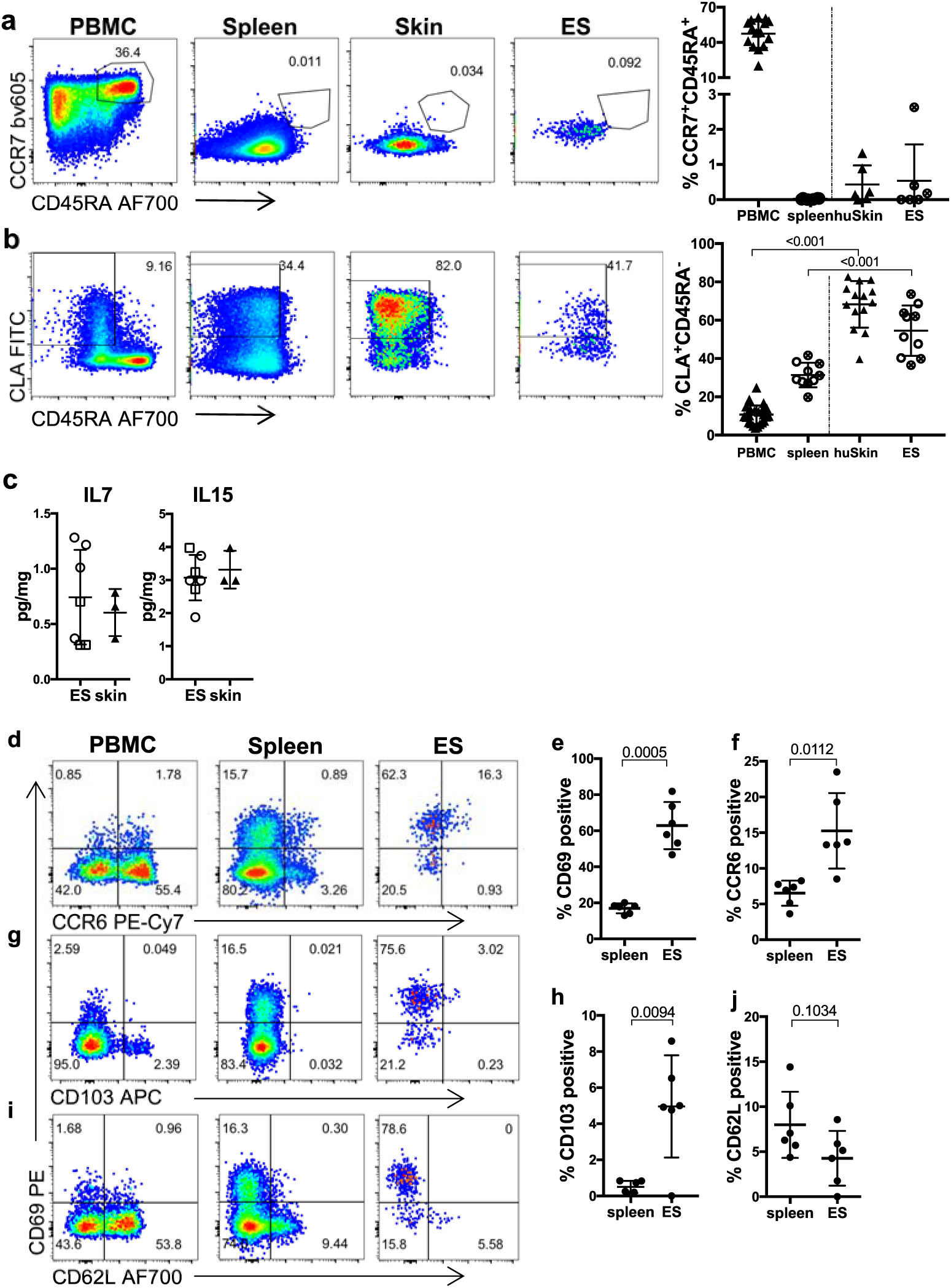
Skin and spleen infiltrating CD4^+^ T cells show skin-homing memory phenotype and upregulate markers of tissue residency and skin-tropism in ES. Representative flow cytometry analysis of (**a**) CCR7 and CD45RA expression, and (**b**) CLA and CD45RA expression by gated CD4^+^CD3^+^CD45^+^ live leukocytes from blood and skin of healthy donors, spleen and ES of huPBMC-ES-NSG mice and graphical summary of the proportions of indicated cells by gated CD4^+^CD3^+^CD45^+^ live leukocytes. n=5-6/experiment; cumulative data of 2 independent experiments. (**c**) Amount of the indicated cytokines per mg skin determined by bead-based multicomponent analysis of ES from huPBMC-ES-NSG and 3 different healthy human skin donors. (**d, g, h**) representative flow cytometry analysis for expression of (**d**) CCR6 and CD69; (**g**) CD62L and (**h**) CD103 in indicated tissues by CLA^+^CD45RA^-^CD4^+^CD3^+^ living cells from (**d**). (**e, f, i, j**) Graphical summary of CLA^+^CD45RA^-^CD4^+^CD3^+^living cells isolated from spleen and ES, expressing the indicated markers. n=5; Significance determined by paired student’s t test; mean ^+^/- SD

### Cutaneous and splenic T cells from huPBMC-ES-NSG mice display multifunctional profiles of T cells in human skin and blood

Next, we sought to determine whether the diverse functional phenotypes of human memory T cells were maintained within the model and thus would be suitable to study human T cell function within human skin *in vivo*. We assessed the function of splenic and ES-derived T cells following *ex vivo* stimulation and intracellular cytokine staining. The ability to produce the Th2, Th17 and Th22 cytokines IL-13, IL-17 and IL-22, respectively, were preserved in CD4^+^ T cells isolated from the huPBMC-ES-NSG mouse when compared to T cells from human blood and skin (Fig. 4 a, b; d-f). By contrast, increased percentages of CD4^+^ T cells isolated from the spleen and ES produced GM-CSF (Fig. 4 c,g). Interestingly while IFNγ^+^CD4^+^ cells were increased in the spleen when compared to PBMC, the proportion of IFNγ-producing CD4^+^ cells within the ES was comparable to skin from healthy donors (Fig. 4 c, h).

**Figure 4:**
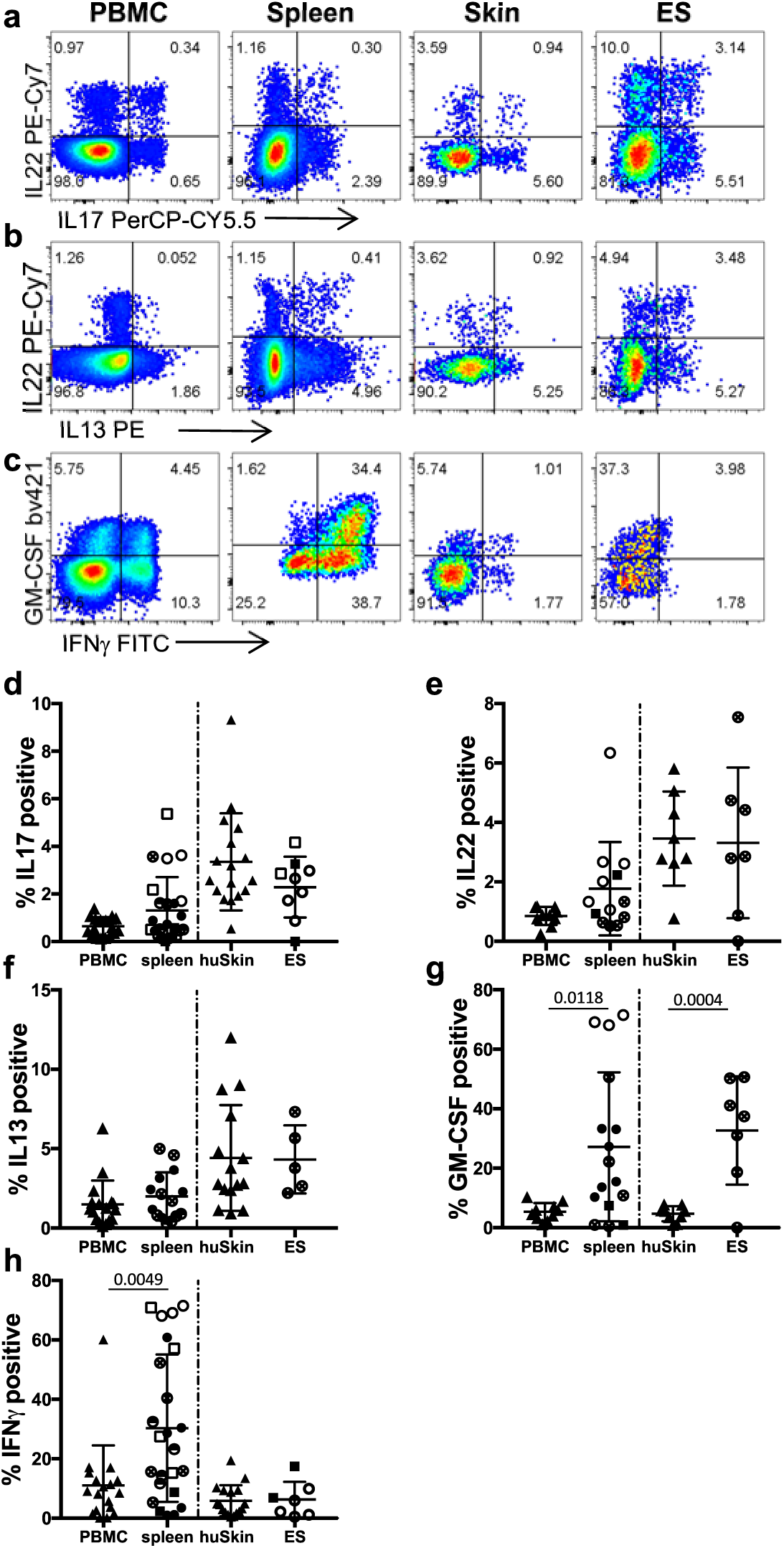
Engrafted splenic and cutaneous human CD4^+^ T cells reflect diverse phenotypes of T cells in human tissues. (**a-c**) Single cell suspensions of blood and skin of healthy donors, and spleen and ES of huPBMC-ES-NSG mice were prepared 18-35 days after PBMC transfer, stimulated *ex vivo* with PMA/ionomycin and intracellular cytokine production was analyzed by flow cytometry. Representative analysis of IL17, IL22, IL13, GM-CSF and IFNγ % of CD4^+^ cells as indicated. (**d-h**) Graphical summary of the expression of the indicated cytokines by T cells from blood and skin of healthy donors and spleen and ES of huPBMC-ES-NSG mice analyzed 18-35 days after PBMC transfer by gated CD4^+^CD3^+^CD45^+^ live leukocytes. n=3-6/experiment; cumulative data of 2-5 independent experiments as indicated by the symbol fillings; each symbol shape is representative of a skin donor (circles = donor WT85 and squares = donor WT70), and each filling represents an independent experiment.

Analogous to the cytokine profiles of splenic and cutaneous CD4^+^ T cells, we assessed the cytokine secretion of CD8^+^ T cells isolated from spleen and ES (Fig. S4). The cytokine profiles of CD8^+^ T cells in ES and spleen were comparable to healthy human skin and PBMC with the exception of GM-CSF (Fig. S4d), which was increased within the ES, similar to the CD4^+^ T cell population. This increased production of GM-CSF might be a result of xenogeneic T cell activation within the model ^55,56^.

As shown before for the memory surface phenotype of T cells within the ES (Fig. 3 and Fig. S3), circulating T cells derived from human blood also assume the functional profile of cutaneous T cells found within human skin upon entry into the ES.

### Cutaneous CD4^+^ T cells are locally activated by microbial antigen

Skin CD4^+^ T cells play a crucial role in controlling cutaneous microbes ^57^. Particularly, the specific role of CD4^+^ T cells in responses against the commensal fungus *Candida albicans* (*C.albicans*) is underscored by the fact that primary and acquired immunodeficiencies that lead to the impairment of CD4^+^ T cell immunity can cause pathogenic *C.albicans* infections ^58–62^. Consistent with that, the human circulating T cell pool contains skin-tropic *C.albicans*-specific memory T cells ^63,64^. Hence we hypothesized that *C.albicans*-specific memory T cells would be present among the adoptively transferred human PBMC and we chose to assess the functionality of these T cells *in vivo*. To evaluate whether a detectable population of *C.albicans*-specific CD4^+^ memory T cells was indeed present in the human PBMC we used for adoptive transfer, we co-cultured donor PBMC for 7 days with autologous monocyte derived dendritic cells (moDCs) that were loaded with heat killed *C.albicans* (HKCA) because *C.albicans* specific T cell responses depend on the presence of HLA-DR^+^ APC ^16,63^. Indeed, we found antigen-specific proliferation, activation and cytokine secretion by HKCA stimulated CD4^+^ T cells when compared to co-cultures of PBMC with non-activated or LPS activated moDCs (Fig. S5 a-c).

Next, we aimed to assess whether this *C.albicans*-specific CD4^+^ memory population would infiltrate the ES and mount a local antigen-specific memory T cell response upon encounter of microbial antigen. However, consistent with previous reports we found poor engraftment of HLA-DR^+^CD3^-^ antigen presenting cells (APC) within the NSG mice ^26^ both in the spleen and ES (Fig. S6). We further found that injection of HKCA alone into the ES had no impact on T cell proliferation or numbers within the ES (Fig. S7). Thus, to compensate for this lack of APC we pulsed autologous moDCs with HKCA (HKCA/moDC) and injected these intradermally into the ES. LPS activated moDC (LPS/moDC) served as a control for non-*C.albicans*-specific activation of T cells by activated APCs. Injections were repeated 3 times within 7 days and ES and spleen were analyzed by flow cytometry one week after the last injection (Fig. 5a). Whereas the proportion of human CD45^+^ cells in the spleen remained unaffected irrespective of the treatment, a slight increase in the percentage of human CD45^+^ cells could be detected in ES injected with HKCA/moDC compared to LPS/moDC injected ES (Fig. 5b). Additionally, a significantly increased proportion of cutaneous CD4^+^ T cells expressed the proliferation marker Ki67 and upregulated CD25 upon injection of HKCA/moDC, indicating activation of CD4^+^ T cells in response to the encountered antigen (Fig. 5c). Similarly, CD4^+^ T cells from ES injected with HKCA/moDC produced significantly higher levels of the effector cytokines IL-17 and TNFα after *ex vivo* PMA/Ionomycin stimulation compared to ES that had received LPS/moDC (Fig. 5d). Importantly, the increased proliferation of CD4^+^ T cells in response to antigen was locally restricted to the injected ES and absent in splenic T cells (Fig. S7c).

**Figure 5:**
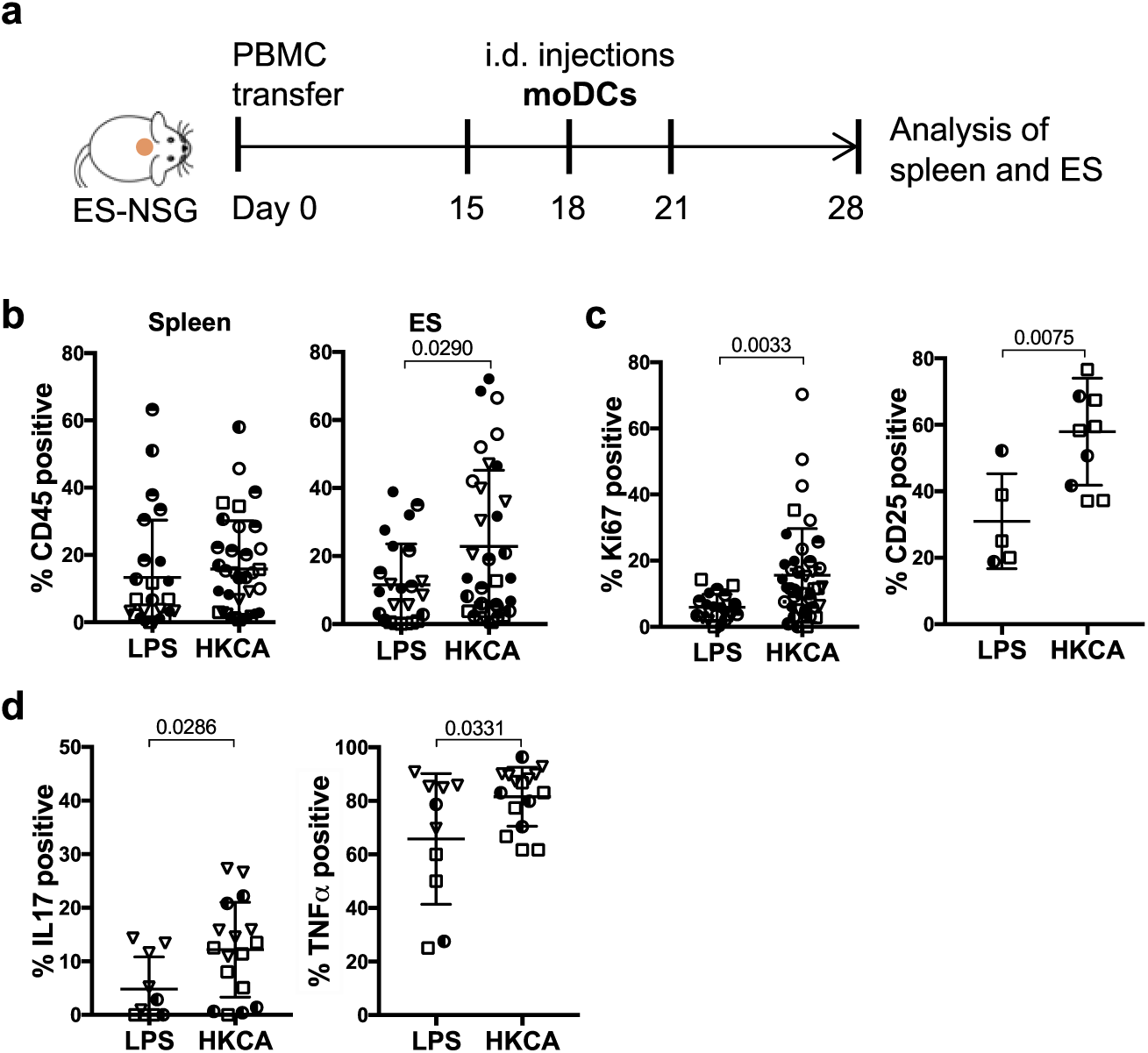
Cutaneous CD4^+^ T cells are activated by local *C.albicans* presented by moDCs in ES. (**a**) Schematic outline of the experiment. (**b**) Graphical summary of the proportion of CD45^+^ cells among live cells in the lymphocyte gate in indicated organs of huPBMC-ES-NSG mice that received either LPS/moDC injections or HKCA/moDC injections into the ES. n=2-7/experiment, cumulative data of 6 independent experiments. (**c**) Graphical analysis of the proportion of Ki67^+^ proliferating cells and CD25^+^ cells by gated CD4^+^CD3^+^CD45^+^ live leukocytes from LPS/moDC or HKCA/moDC treated ES. (**c**) Single cell suspensions of ES were analyzed by flow cytometry after *ex vivo* stimulation with PMA/Ionomycin and intracellular cytokine staining. Graphical summary of the proportion of skin CD4^+^ T cells by gated CD4^+^CD3^+^CD45^+^ live leukocytes expressing IL17 and TNFα. n=2-7/experiment, cumulative data of 3 independent experiments. (circles = donor WT85, squares = donor WT70, triangles = WT73) Statistical significance determined by 2-tailed unpaired student’s t test; mean +/- SD.

### Antigen-specific T cell responses remain detectable in donor-mismatched skin tissue

These initial experiments were all performed using a completely matched system where ES, PBMC and moDC were from the same donor. However, access to skin that is matched to the PBMC of a specific patient group may present a limiting factor in studies of human cutaneous immune responses. To broaden the model’s applicability, we sought to determine whether antigen-specific T cell responses could also be detected in the ES when we used donor-mismatched tissues. Therefore, we compared cutaneous CD4^+^ T cell responses to HKCA in donor-matched and -mismatched ES. ES were generated from two different donors (donor A or B) designated ES-NSG-A and ES-NSG-B. After complete wound healing, both recipients received PBMC that were either matched or mismatched to the ES and were injected intradermally with matched LPS/moDC or HKCA/moDC (i.e. the leukocyte populations were always HLA-matched) (Fig. 6a). The experiments were performed in both directions with A and B being either ES or leukocyte donor.

**Figure 6:**
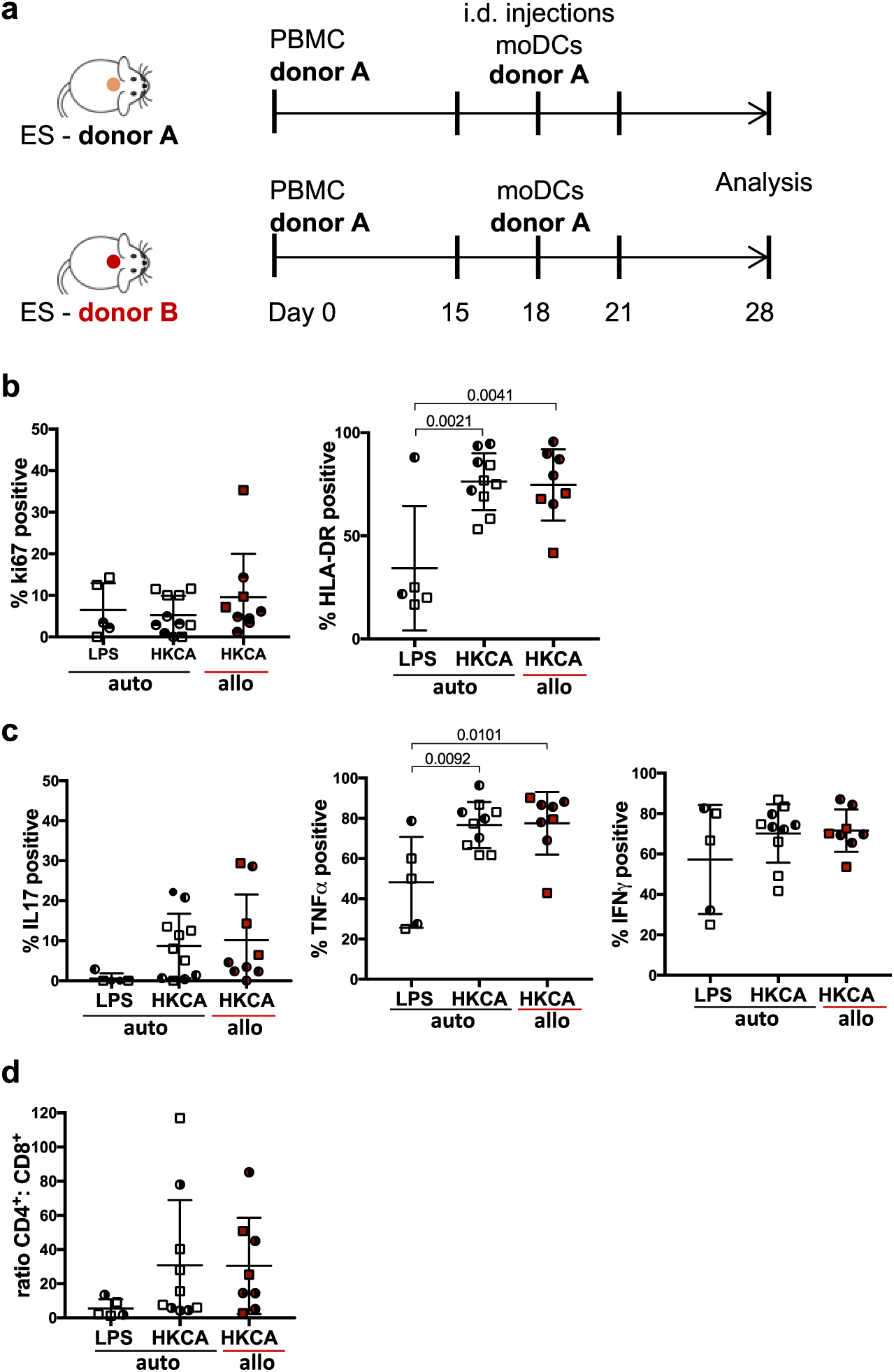
*C.albicans*-specific CD4^+^ T cell response can be detected in allogeneic ES. (**a**) Schematic: NSG mice bearing fully healed ES of one of two different skin donors (A and B) were adoptively transferred with either skin donor-matched PBMC or skin donor-mismatched PBMC. Intradermal injections of donor A derived LPS/moDC or HKCA/moDC were performed as depicted (i.e. leukocytes were matched). Single cell suspensions of ES were analyzed by flow cytometry after *ex vivo* stimulation with PMA/Ionomycin and intracellular staining. (**b**,**c**) Graphical summary of the proportion of skin CD4^+^ T cells by gated CD4^+^CD3^+^CD45^+^ live leukocytes expressing the indicated markers following intradermal encounter of LPS/moDC (LPS) or HKCA/moDC (HKCA). Red data points represent CD4^+^ T cells isolated out of mismatched ES. Statistical significance determined by ANOVA and Tuckey’s test for multiple comparison; mean +/-SD. (**d**) Graphical summary of the ratio between CD4^+^ and CD8^+^ T cells of isolated skin T cells gated by CD3^+^CD45^+^ live leukocytes. n=2-5/group, combined data of 2 independent experiments;

The proliferation of CD4^+^ T cells within allogeneic ES was slightly but not significantly increased compared to the fully matched system (Fig. 6b). Additionally, the levels of the activation maker HLA-DR ^65–67^ were comparable within the allogeneic ES tissue compared to the autologous ES in response to HKCA/moDC, and significantly increased compared to LPS/moDC injected ES (Fig. 6b). Similarly, CD4^+^ T cells within the ES injected with HKCA/moDC secreted IL17 and TNFα compared to LPS/moDC, indicating *C.albicans*-specific activation of the T cells (Fig. 6c). Importantly, similar to HLA-DR the proportion of IFNγ^+^ CD4^+^ T cells were not increased within the allogeneic ES (Fig. 6c). Additionally, CD4:CD8 ratios remained unchanged between skin T cells from matched and mismatched HKCA/moDC injected ES suggesting a lack of CD8 expansion in response to the allogeneic keratinocytes and fibroblasts (Fig. 6d). Splenic CD4^+^ T cells showed no indication of an allogeneic response or HKCA-specific cytokine production (Fig. S8 a-d) and, splenic CD4^+^:CD8^+^ T cell ratios were unaltered in response to the allogeneic ES (Fig. S8e), indicating the absence of a systemic response.

### CD4^+^ T cells isolated from ES respond to HKCA antigen *ex vivo*

To further confirm that the local activation and cytokine response of cutaneous T cells was truly antigen-specific rather than a non-specific response that was promoted by PMA/ionomycin stimulation, we isolated cutaneous T cells from ES that had been injected with HKCA/moDC, and then restimulated these cells ex vivo with moDCs that were either activated with LPS or loaded with HKCA in the presence of Brefeldin A (Fig. 7a). We found that re-stimulation with HKCA/moDCs resulted in an increased fraction of proliferating Ki67 positive CD4^+^ T cells (Fig.7b) and increased effector cytokine production compared to T cells that were re-stimulated with LPS/moDCs (Fig.7c). These results indicate that the increased effector response of cutaneous T cells (observed in Figs. 5 and 6) is due to antigen-specific activation. Furthermore, we found that the injection of free HKCA (without moDCs) into the ES did not induce increased immune cell numbers or proliferation (read out as KI67^+^ proportions), compared to the rate of CD4^+^ T cells that were stimulated with LPS/moDC (Fig. S7). This indicates that the described response of CD4^+^ T cells in the HKCA/moDC group is dependent on antigen-presentation and not due to pathogen-associated signals present in HKCA.

**Figure 7:**
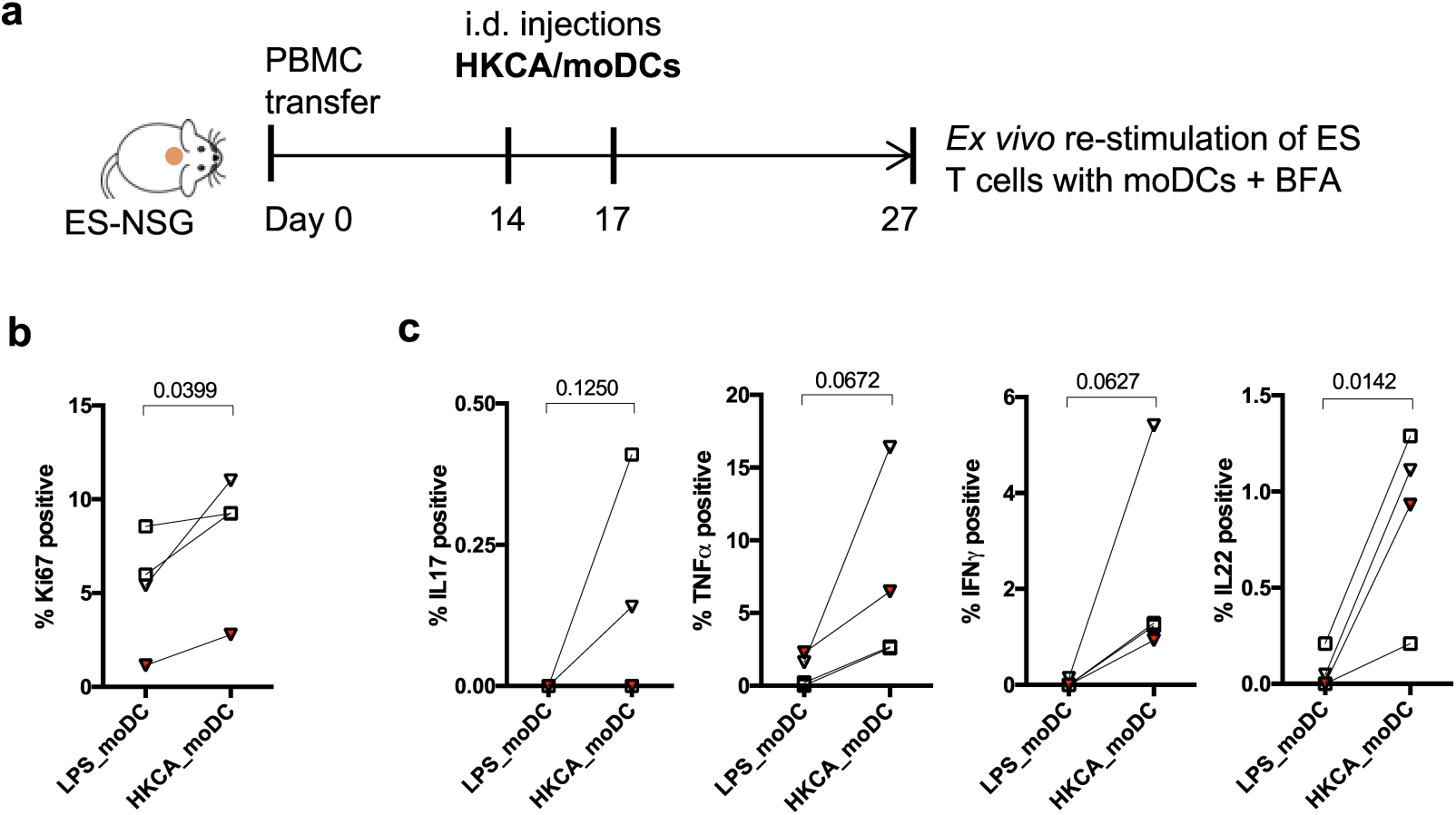
CD4^+^ T cell activation within the *C.albicans* injected ES grafts is antigen-specific. (**a**) Schematic of experimental procedure. Mice carrying ES were transferred with hPBMC followed by 1-2 intradermal injections of HKCA*-*loaded moDCs autologous to the PBMC. 10 days after the last injection, single cell suspensions of ES were divided and re-stimulated *ex vivo* with autologous moDCs stimulated with LPS or loaded with HKCA in presence of Brefeldin A (BFA) to detect cytokines by flow cytometry. (**b**) Graphical analysis of the proportion of Ki67^+^ proliferating cells of gated CD4^+^CD3^+^CD45^+^ live leukocytes from ES re-stimulated with LPS/moDC or HKCA/moDC. (**c**) Graphical summary of the proportion of CD4^+^ T cells from ES re-stimulated with LPS/moDC or HKCA/moDC producing the indicated cytokines. Combined data of 3 independent experiments (n=3-11 mice/experiment). Squares = donor WT70, triangles = WT73. Red data points represent CD4^+^ T cells isolated out of mismatched ES. Clear symbols indicate autologous setting. Statistical significance determined by one tailed paired student’s t test.

### Increased effector function of CD69^+^ CD4^+^ T_RM_-like T cells in response to *C.albicans*

In a recent study it was shown that *C.albicans* specific responses in murine skin was mostly mediated by a population of cutaneous CD69^+^CD4^+^ T_RM_ cells generated after primary *C.albicans* infection, and a similar population was isolated from human skin ^16^. Importantly, we identified a CD69^+^ population present within the ES prior to the encounter of *C.albicans* within the *ES* (Fig. 3e). We hypothesized that these newly established CD69^+^ CD4^+^ memory-type T cells would show increased effector function upon antigenic challenge with the skin microbe *C.albicans* compared to the CD69^-^ population, similar to what has been shown in human skin ^16^. In line with these *ex vivo* findings, the CD69^+^CD4^+^ T cell population isolated from ES that has been injected with HKCA/moDC showed superior effector functions compared to the CD69^-^ counterpart (Fig. 8). It remains to be determined whether this CD69^+^ CD4^+^ T_RM_-like cell population was recruited upon application of HKCA/moDC or seeded prior to *C.albicans* injection from the circulating pool of *C.albicans* specific T cells.

**Figure 8:**
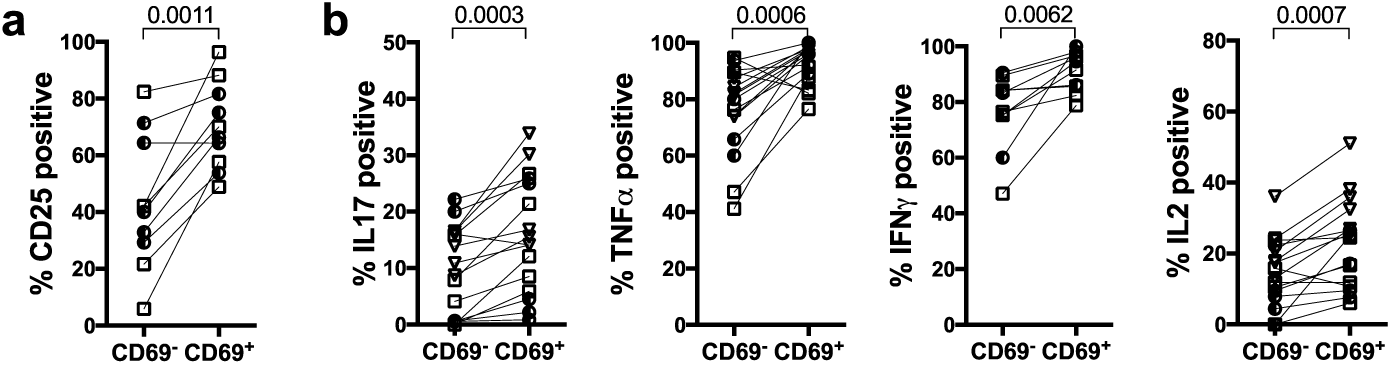
Cutaneous CD4^+^CD69^+^ T Cells show increased effector function in response to *C.albicans*. Mice were treated as in Fig.5. 7 days after the last injection of HKCA/moDC single cell suspensions were stimulated with PMA/ionomycin and analyzed by flow cytometry Graphical summary of the expression of (**a**) CD25 or (**b**) the indicated cytokines by CD69^-^ or CD69^+^ gated CD4^+^CD3^+^CD45^+^ live leukocytes from ES. Each symbol represents an individual animal, donors: circles = donor WT85, squares = donor WT70, triangles = WT73; data from 2-3 experiments. Significance determined by paired student’s t test.

## Discussion

Human skin contains a significant number of memory T cells that provide protective immunity and support tissue homeostasis ^2,3,5,25,35^. The generation of cutaneous resident T_RM_ cells at the site of primary infection has been studied using murine models ^6,7^. However, suitable *in vivo* models to study human memory T cell generation, migration, and function of human cutaneous T cells to promote translational research were still lacking, and existing skin-humanized mouse models almost exclusively use allogeneic or inflammatory settings ^25,28,30,31^. Fundamental insights revealing the heterogeneity of cutaneous human memory T cell subsets were gained recently by the group of Rachel Clark, when they investigated memory T cell populations in human skin in a xenograft skin mouse model. However, the infiltration of PBMC into the skin was driven by allogeneic MHC recognition of donor APCs contained in the skin graft ^25^, thus reflected an inflammatory skin condition. To avoid the presence of resident immune cells within skin humanized mouse models, bioengineered skin combined with intradermal injection of *in vitro* generated T cell subsets and recombinant cytokines into the skin graft was used to study the pathogenesis of atopic dermatitis (AD) and psoriasis ^30,31^. However, these models did not follow cutaneous functions of skin-tropic T cells (i.e. a memory sub-population specialized in cutaneous immunity) ^68–70^. In health and disease, T cell recruitment to and function within the skin depends on a variety of skin-derived chemokines and cytokines ^5^, and mutations leading to the loss of signaling result in impaired cutaneous T cell recruitment and/or maintenance ^38,51,52^. Importantly, the ES we used ^41^ reflected the chemokine environment found in healthy human skin, particularly chemokines and cytokines involved in T cell recruitment and activation, such as CCL2 ^46^, CCL5 ^71^, CXCL10 ^48^, CXCL12 ^49^ and T cell function and maintenance, like IL7 ^51,52^ and IL15 ^51,53^. In line with this, T cells migrated into the ES upon adoptive transfer of human PBMC, a process that was likely not driven by a tissue damage response, because the levels of inflammatory cytokines such as IL1α, IL1β, TNFα, TNFβ, IL18 and IL23 were lower or at equal levels in the ES when compared to healthy human skin. This highlights the power of the huPBMC-ES-NSG model to allow the study of human memory T cell migration and function in absence of acute inflammation, as well as the impact of tissue-derived signals on immunological processes in the skin. Importantly, the use of ES (i.e. human skin tissue without passenger leukocytes) permits precise control over the cell populations that partake in a specific immune response.

The ES was infiltrated by diverse subsets of memory T cells that maintained the multifunctional profiles of T cells found in human skin, and this was independent of the presence of APCs within the model. Importantly, T cells within the ES were functionally remarkably similar to T cells isolated from human skin, underlining the applicability of the huPBMC-ES-NSG model in studies of cutaneous T cell function *in vivo*. However, due to the increased proportion of GM-CSF secreting T cells in the system, which is likely the result of xenogeneic T cell activation within the model ^55,56^, we expect that GM-CSF production would be a poor read-out for antigen-specific immune responses (e.g. against *C.albicans*). Thus, while GM-SCF plays an important role in anti-microbial immune responses ^56,72^, we believe that our skin-humanized mouse model is not suitable to study the function and regulation of GM-CSF.’

As in human subjects ^2^, blood-derived skin-tropic CLA^+^CD4^+^ and CD8^+^ T cells accumulated within the skin when compared to splenic T cells and increased their expression of T_RM_ markers, such as CD69 and CD103 ^15,25^, underscoring the unique potential of this model to study the impact of tissue-derived factors on memory T cell generation.

It has been previously shown that in patients that had received intestinal allografts, recipient-derived graft-infiltrating CD69^-^ T cells gave rise to long-lasting CD69^+^ T_RM_ that replenished the total gut-resident T_RM_ compartment ^73^. A similar process of *de novo* T_RM_ generation has been observed in patients that had received allogeneic lung transplants ^8^. In line with that, the generation of CD69^+^ T_RM_-like cells within the ES occurred from a pool of circulating CD69^-^ T cells in absence of increased inflammatory cues or microbial antigen presentation within the ES.

Consistent with a population of *C.albicans*-specific cutaneous CD69^+^CD4^+^ T_RM_ cells isolated from human skin ^16^, CD69^+^CD4^+^ T cells responded more vigorously to injection of HKCA/moDCs into the ES. In fact, superior effector function of CD69^+^ T_RM_-like T cells has recently been shown for human T_RM_ in various tissue sites ^21^ when compared to the CD69^-^ CD4^+^ T cells. It remains to be determined whether this CD69^+^ CD4^+^ T_RM_-like cell population was seeded prior to *C.albicans* injection from the circulating pool of *C.albicans* specific T cells. The fact that the model closely reflects the immunological response towards *C.albicans* makes it a formidable tool to study the dynamics and requirements of human cutaneous T_RM_ function *in vivo*, and potentially facilitating translational research in T_RM_-mediated diseases.

We found that intradermal injection of moDCs loaded with antigen could substitute for poor engraftment of APC within the NSG model. Interestingly, antigen-specific T cell activation in response to HKCA presented by matched moDCs was detectable in both, autologous and allogeneic ES. Thus, access to matched blood and tissue samples might not be limiting the study of cutaneous CD4^+^ T cell responses in these skin-humanized mice.

Together, these data suggest that the huPBMC-ES-NSG model represents a suitable tool to study *C.albicans* specific local activation and memory responses of cutaneous T cells *in vivo* in a non-inflammatory setting. Importantly, the keratinocytes and fibroblasts used to generate ES can be cultured and manipulated using techniques such as CRISPR technology^74^. Thus, the huPBMC-ES-NSG model provides a highly versatile tool to study cutaneous T cell responses and to manipulate tissue-derived signals that impact skin immunity. Additionally, the model may serve as a platform to test novel therapeutic interventions to treat cutaneous inflammation, skin tumors or autoimmune diseases.

## Material and Methods

All methods were carried out in accordance with the relevant guidelines and regulations.

### Mice

All animal studies were approved by the Austrian Federal Ministry of Science, Research and Economy. NOD.Cg-Prkdcscid Il2rgtm1Wjl/SzJ (NSG) mice were obtained from The Jackson Laboratory and bred and maintained in a specific pathogen-free facility in accordance with the guidelines of the Central Animal Facility of the University of Salzburg.

### Human specimens

Normal human skin was obtained from patients undergoing elective surgery, in which skin was discarded as a routine procedure. Blood and/or discarded healthy skin was collected at the University Hospital Salzburg, Austria. Informed consent was obtained from all subjects. Samples of subjects of both sexes were included in the study. All studies were approved by the Salzburg state Ethics Commission (decision: according to Salzburg state hospital law no approval required) (Salzburg, Austria).

### PBMC isolation for adoptive transfer into NSG recipients and flow cytometry

Human PBMC were isolated from full blood using Ficoll-Hypaque (GE-Healthcare; GE17-1440-02) gradient separation. PBMC were frozen in FBS with 10% DMSO (Sigma-Aldrich; D2650), and before adoptive transfer thawed and rested overnight at 37°C and 5% CO_2_ in RPMIc (RPMI 1640 (Gibco; 31870074) with 5% human serum (Sigma-Aldrich; H5667 or H4522), 1% penicillin/streptomycin (Sigma-Aldrich; P0781), 1% L-Glutamine (Gibco; A2916801), 1% NEAA (Gibco; 11140035), 1% Sodium-Pyruvate (Sigma-Aldrich; S8636) and 0.1% β-Mercaptoethanol (Gibco; 31350-010). Cells were washed and 1.8-3×10^6^ PBMC/mouse intravenously injected. Female and male donors for adoptive transfer into huPBMC-NSG mice were aged 40-55 years.

Murine neutrophils were depleted with mLy6G (Gr-1) antibody (BioXcell; BE0075) intraperitoneally every 5-7 days as described before ^28^.

### Generation of engineered skin (ES)

Human keratinocytes and fibroblasts were isolated from human skin and immortalized using human papilloma viral oncogenes E6/E7 HPV as previously described ^40^. For all experiments two different skin donors were used: WT70 (indicated with square symbols in graphs), and WT85 (round symbols). Cells were cultured in Epilife (Gibco, MEPICF500) or DMEM (Gibco; 11960-044) containing 2% L-Glutamine, 1% Pen/Strep, 10% FBS, respectively. Per mouse, 1-2×10^6^ keratinocytes were mixed 1:1 with autologous fibroblasts in 400µl MEM (Gibco; 11380037) containing 1% FBS, 1% L-Glutamine and 1% NEAA for *in vivo* generation of engineered skin as described ^41^, with slight variations, specifically the use of immortalized keratinocytes and fibroblasts. Additionally, the silicone grafting chambers were removed completely 7 days after transplantation.

### T cell isolation from skin tissues for flow cytometry

Healthy human skin and ES were digested as previously described ^29^. Approximately 1cm^2^ of skin was digested overnight in 5%CO_2_ at 37°C with 3ml of digestion mix containing 0.8mg/ml Collagenase Type 4 (Worthington; #LS004186) and 0.02mg/ml DNase (Sigma-Aldrich; DN25) in RPMIc. ES were digested in 1ml of digestion mix. Samples were filtered, washed and stained for flow cytometry or stimulated for intracellular cytokine staining. Approx. 3 cm^2^ of shaved dorsal mouse skin were harvested and single cell suspensions prepared as described ^78^ and stained for flow cytometry.

### Generation of monocyte derived dendritic cells (moDC)

moDC were generated from frozen PBMC similar to what has been described previously ^79^. Briefly, PBMC were thawed and monocytes adhered for 75 min at 37 °C and 5% CO_2_ in DC medium (RPMI 1640: 10% FBS, 2 mM L-Glutamine, 100 U/ml penicillin/streptomycin, 50 μM β-mercaptoethanol). After washing, adherent monocytes were cultured in DC medium supplemented with 50 ng/ml GM-CSF (ImmunoTools; 11343127) and 50 ng/ml IL4 (ImmunoTools; 11340047) for 7 days to generate immature DC. After 6 days, cells were harvested and re-plated in DC medium without cytokines. For activation, moDCs were cultured for 9-13 hrs with 5ng/ml LPS (Sigma-Aldrich; L2880) or 10^6^ cells/ml heat killed *Candida albicans* (eubio; tlrl-hkca). 1.8-3×10^4^ moDC/mouse were intradermally injected in 50µl PBS/mouse.

### Antigen-specific re-stimulation of skin-derived T cells

Single cell suspension of ES pooled from multiple mice were divided into two equal parts and stimulated with LPS or HKCA loaded moDC autologous to the T cells for 20 hrs, 5µg/ml Brefeldin A was added after 1 hr of stimulation.

### Flow cytometry

Cells were stained in PBS for surface markers. For detection of intracellular cytokine production, spleen and skin single cell suspensions and PBMC were stimulated with 50 ng/ml PMA (Sigma-Aldrich; P8139) and 1 µg/ml Ionomycin (Sigma-Aldrich; I06434) with 10 µg/ml Brefeldin A (Sigma-Aldrich; B6542) for 3.5 hrs. For permeabilization and fixation Cytofix/Cytoperm (BectonDickinson; RUO 554714) or Foxp3 staining kit (Invitrogen; 00-5523-00) were used. Data were acquired on LSR Fortessa (BD Biosciences) or Cytoflex LS (Beckman.Coulter) flow cytometers and analyzed using FlowJo software (Tree Star, Inc.) A detailed list of the used antibodies can be found in Supplementary Table S1.

### Histological staining of skin sections

Normal human skin, ES and adjacent murine skin were frozen in TissueTek (Sakura; TTEK). 7 µm cryosections were stained with Hemalum solution acid (Carl Rorth; T865.1) and Eosin Y aqueous solution (Sigma, 201192A). Human type VII collagen was stained by immunofluorescence using anti-human type VII collagen antibody and goat anti-rabbit A488 as secondary antibody, ProLong™ Gold Antifade Mountant with DAPI, (Invitrogen; P36931) was used for nuclear staining and mounting. For immunohistochemistry paraffin-embedded normal human skin and ES was stained for human Cytokeratin 5/6 according to the manufacturer’s protocol using a Ventana BenchMark Series automated slide stainer with ultraView Universal DAB Detection kit (Roche, 760-500).

### ProcartaPlex™ immunoassays from human skin and engineered skin

Human skin or ES from huPBMC-ES-NSG mice were stored at −70°C until use. Skin was taken up in PBS with Protease Inhibitor Cocktail (1:100) (Sigma-Aldrich; P8340), homogenized and filtered through 0.22µm SpinX columns. Suspensions were stored at −70°C until use. Samples were used at 8mg/ml for assay. ProcartaPlex immunoassay was performed according to the manufacturer’s protocol and measured using Luminex Magpix® system.

### A detailed list of antibodies and reagents can be found in supplementary Table S1

#### Statistical analysis

Statistical significance was calculated with Prism 7.0 software (GraphPad) by one-way ANOVA with Tukey’s multiple comparisons test, or by paired or unpaired student’s t-test as indicated. Error bars indicate mean ^+^/- standard deviation. Statistical analyses were performed for all data sets and p-values indicated in the graph only when significant changes were observed.

## Acknowledgements

We especially thank all human subjects for blood and skin donation and the nurses at the Breast Center University Hospital of the Paracelsus Medical University Salzburg, Austria. We thank Dr. Stefan Hainzl, EB House Austria, Department of Dermatology, University Hospital of the Paracelsus Medical University Salzburg, Austria, for the immortalization of primary human keratinocytes and fibroblasts. We thank Monika Prinz from the Department of Dermatology at the University Hospital of the Paracelsus Medical University Salzburg, Austria for help with the IHC staining and Peter Steinbacher from the Department of Biosciences at the University of Salzburg, Austria, for support with microscopy. This work was supported by the Focus Program “ACBN” of the University of Salzburg, Austria, by a grant from the Dystrophic Epidermolysis Bullosa Research Association (DEBRA) International and DEBRA Austria, and NIH grant R01AI127726. MMK is part of the PhD program Immunity in Cancer and Allergy, funded by the Austrian Science Fund (FWF, grant W 1213) and was recipient of a DOC Fellowship of the Austrian Academy of Sciences.

## Author Contributions

IKG, EMM, MDR, DJC and MMK conceptualized the study, MMK, EMM and IKG designed the experiments; MMK, AB, LMG, SRV and RH acquired the data; ML performed IHC staining, ASt, RR, and AS acquired human samples, TN and JH-H helped establish the moDC cultures and performed quality control; MMK performed data analysis, MMK and IKG interpreted the data and wrote the manuscript. All authors reviewed the final version of the manuscript. IKG and EMM supervised the project.

## Supplementary data

**Supplementary Figure 1:**
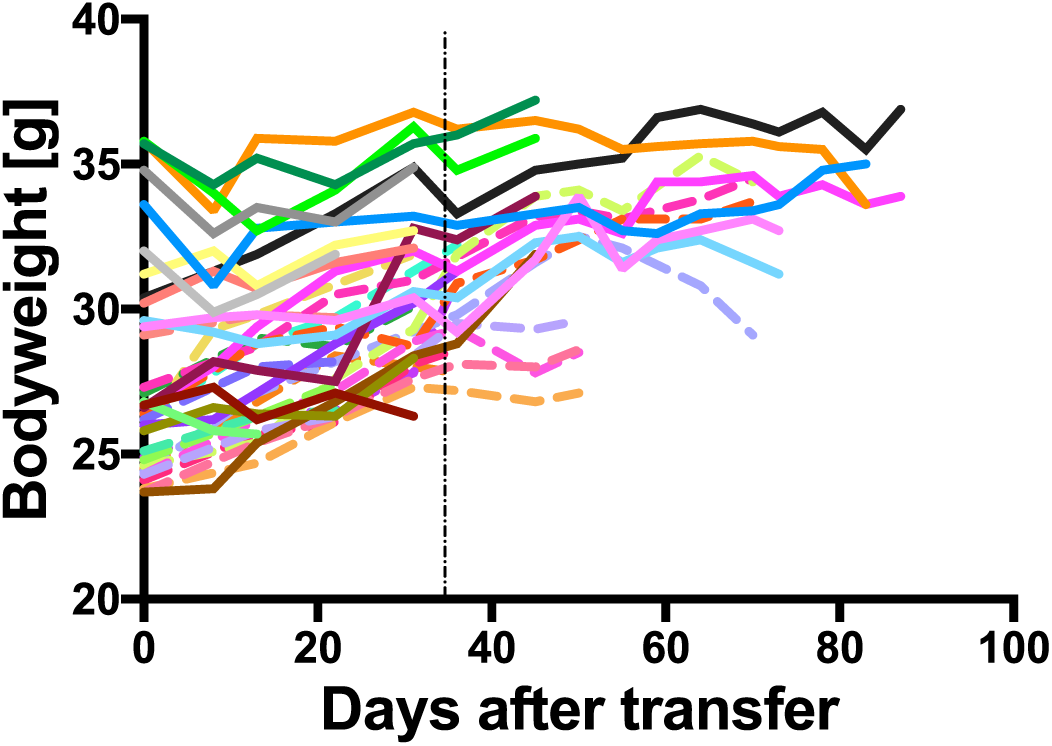
Body weight curves of experimental mice after adoptive transfer human PBMC. NSG mice were adoptively transferred with human PBMC and weight was measured at indicated time points. Graph shows 2 representative experiments of 2 different PBMC donors (full lines and dotted lines) after adoptive transfer of 2.5×10^6^ PBMC; n=36. Vertical black dotted line represents the limit of 35 days for all further experiments using huPBMC-ES-NSG mice.

**Supplementary Figure 2:**
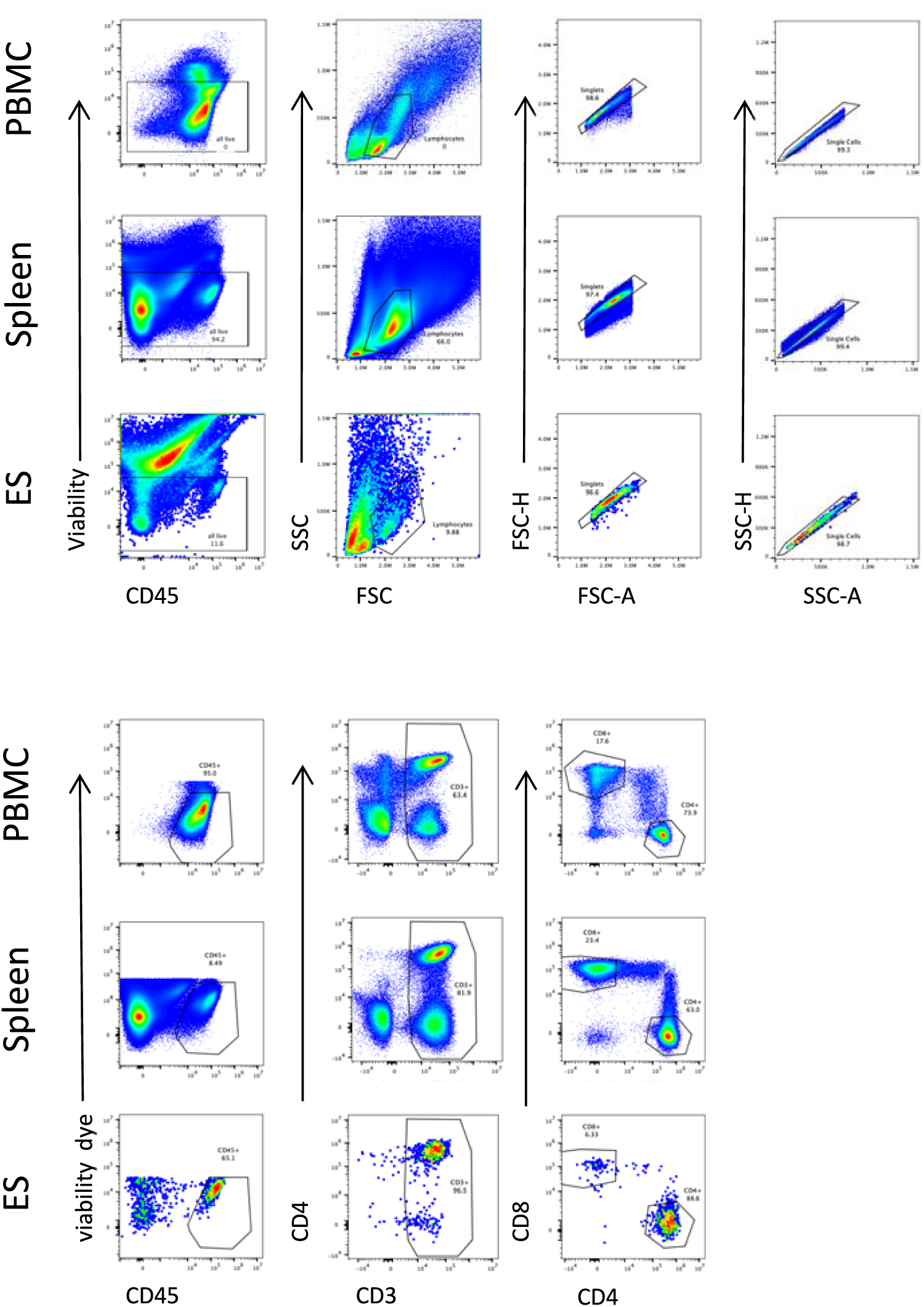
Gating strategy for flow cytometric analysis of human PBMC, spleen and ES single-cell suspensions.

**Supplementary Figure 3:**
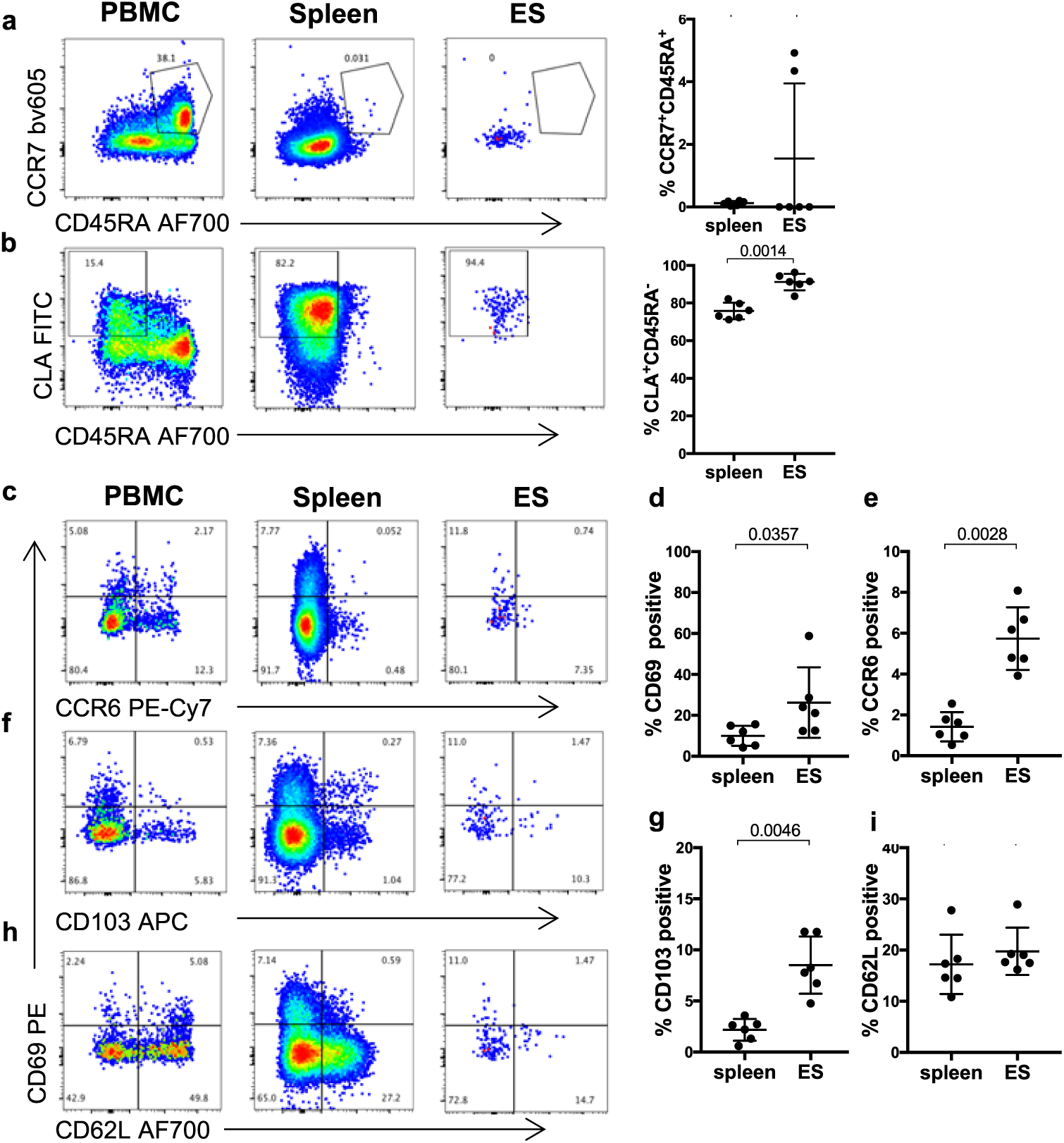
Skin and spleen infiltrating CD8^+^ T cells show skin-homing memory phenotype and upregulate markers of tissue residency and skin-tropism in human ES. Representative flow cytometry analysis of (**a**) CCR7 and CD45RA expression, and (**b**) CLA and CD45RA expression by gated CD4^+^CD3^+^CD45^+^ live leukocytes from blood of healthy donors, spleen and ES of huPBMC-ES-NSG mice and graphical summary of the proportions of indicated cells by gated CD4^+^CD3^+^CD45^+^ live leukocytes. n=5-6/experiment; cumulative data of 2 independent experiments. (**c, f, h**) Representative flow cytometry analysis for expression of (**c**) CCR6 and CD69; (**f**) CD103 and (**h**) CD62L in indicated tissues by gated CLA^+^CD45RA^-^CD4^+^CD3^+^ live cells from blood of healthy donors, spleen and ES of huPBMC-ES-NSG mice. (**d, e, g, i**) Graphical summary of the expression of the indicated markers by CLA^+^CD45RA^-^CD4^+^CD3^+^living cells isolated from spleen and ES. Each dot represents an individual animal; Significance determined by paired student’s t test; mean ^+^/- SD

**Supplementary Figure 4:**
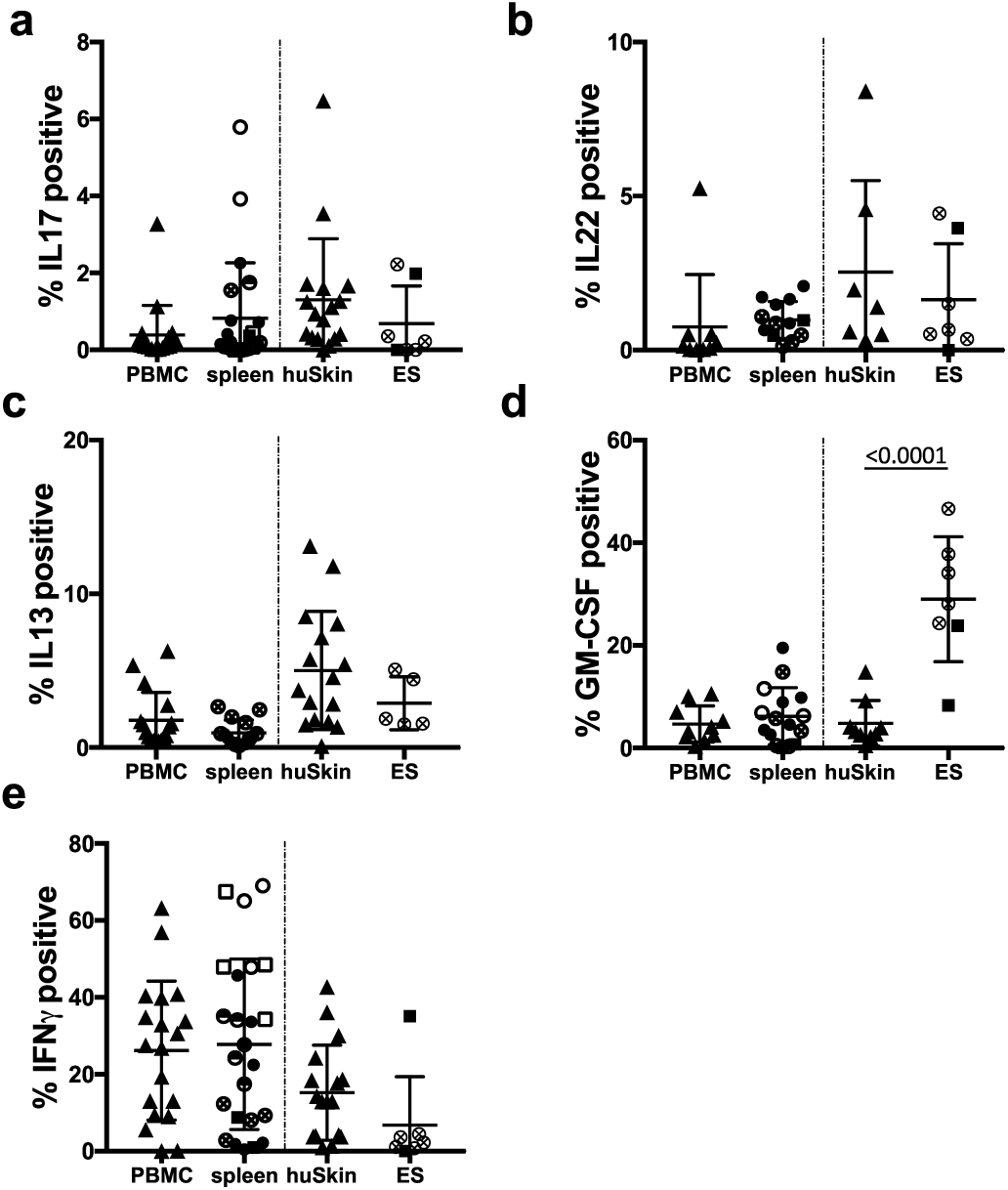
Engrafted splenic and cutaneous human CD8^+^ T cells reflect diverse phenotypes of T cells in human tissues. Single cell suspensions of blood and skin of healthy donors, and spleen and ES of huPBMC-ES-NSG mice were prepared 18-35 days after PBMC transfer, stimulated *ex vivo* with PMA/ionomycin and intracellular cytokine production was analyzed by flow cytometry. (**a-e**) Graphical summary of flow cytometry analysis of IL17, IL22, IL13, GM-CSF and IFNγ producing CD8^+^CD3^+^CD45^+^ gated live leukocytes from blood and skin of healthy donors and spleen and ES of huPBMC-ES-NSG mice as indicated upon *ex vivo* stimulation with PMA/Ionomycin and intracellular staining. n=3-6/ experiment; combined data of 1-4 independent experiments, as indicated by the different fillings of the symbols.

**Supplementary Figure 5:**
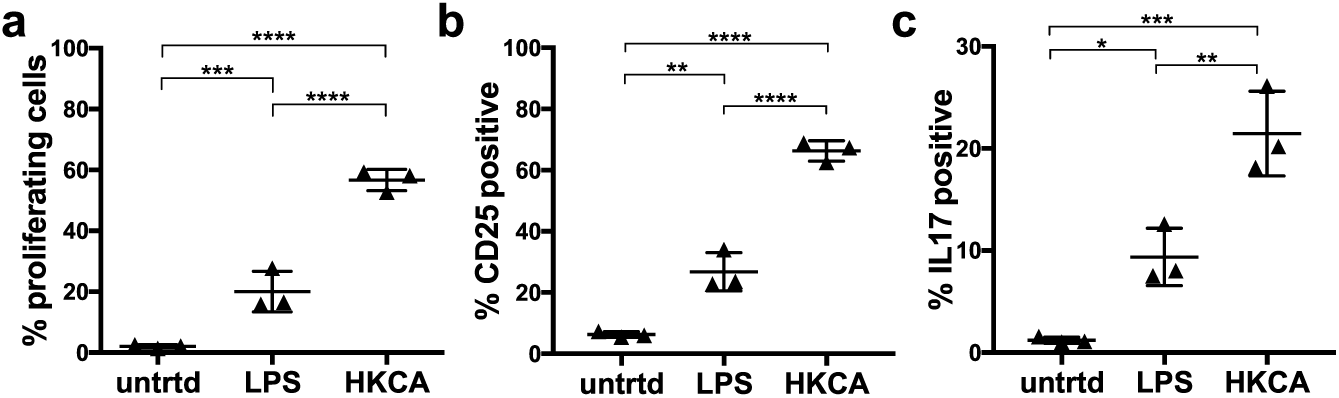
Heat killed *C.alibicans* presented by moDCs leads to proliferation and activation of CD4^+^ T cells. PBMC were cultured *in vitro i*n the presence of monocyte-derived DCs (moDC) for 7 days and the proliferation, CD25 expression and cytokine production of CD4^+^ T cells gated by CD3^+^CD45^+^live leukocytes was assessed by flow cytometry. (**a**) Graphical summary of proliferating CD4^+^ T cells and expression of the indicated markers by of human PBMC with autologous non-activated moDC (untrd), LPS loaded moDC (LPS) or HKCA loaded moDC (HKCA). PBMC were stained with the cell proliferation dye eFluor 450 to identify proliferating cells. n=3/group. P values indicated as asterisk: * ≤ 0.05; ** ≤ 0.01, *** ≤ 0.001, **** ≤ 0.0001

**Supplementary Figure 6:**
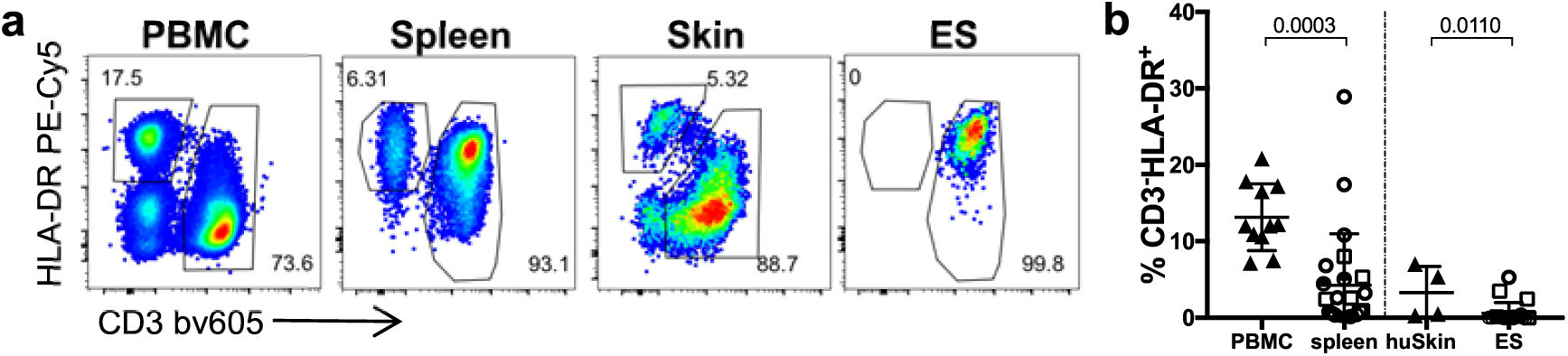
CD3^-^HLA-DR^+^ antigen presenting cells do not engraft well within huPBMC-ES-NSG. **a**) Representative flow cytometry analysis of CD3^-^HLA-DR^+^ cells in human PBMC, skin and spleen and ES of huPBMC-ES-NSG and (**b**) graphical summary of CD3^-^HLA-DR^+^ cells gated on live CD45^+^ leukocytes. n=4-6/group, combined data of 1-5 independent experiments, as indicated by the fillings of the symbols.

**Supplementary Figure 7:**
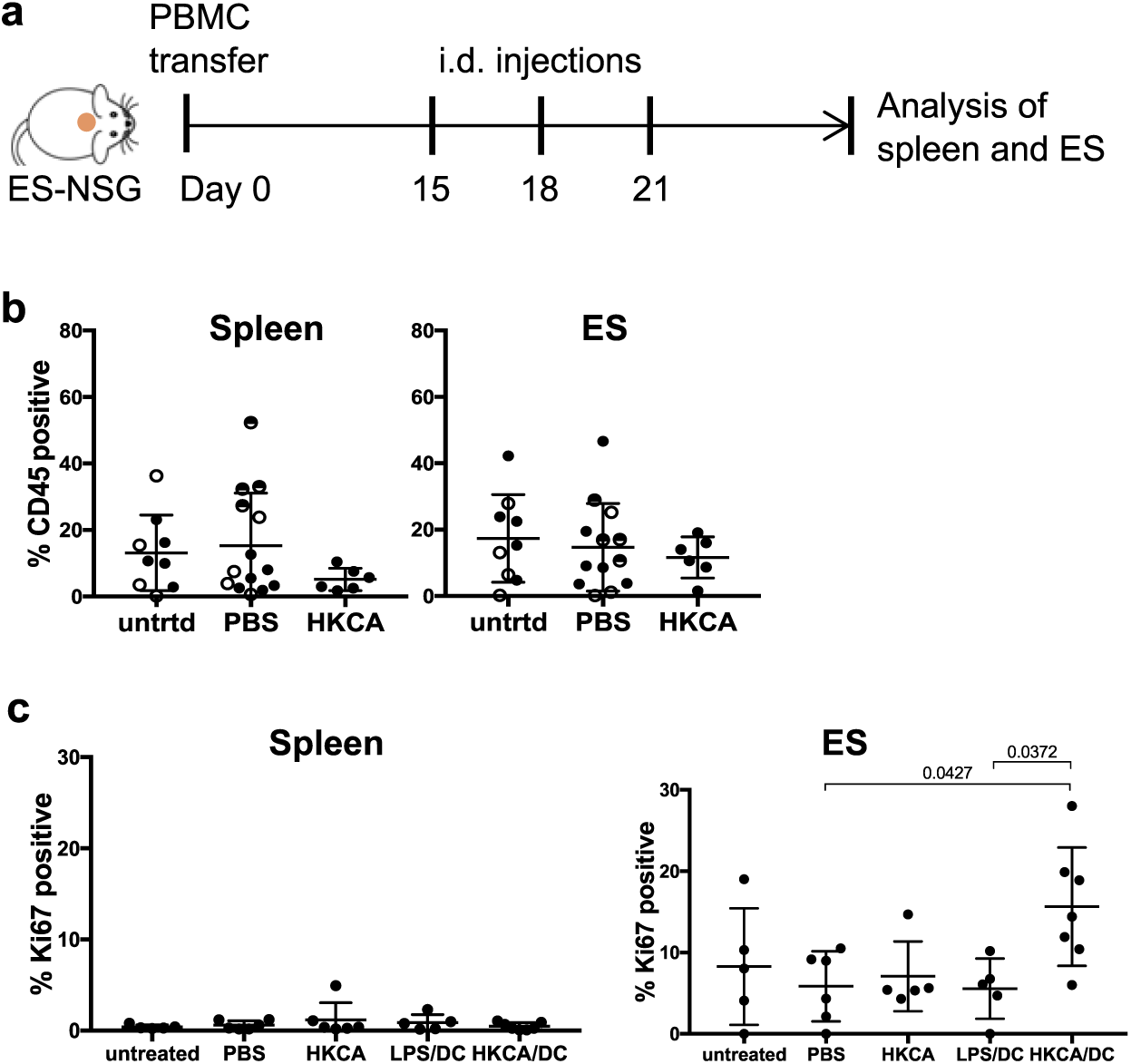
Free heat killed *C.alibicans* fails to elicit proliferation and activation of CD4^+^ T cells. (**a**) Experimental procedure of the injection of 1×10^6^/ml free HKCA cells into the ES. Single cell suspensions of indicated organs were analyzed by flow cytometry. (**b**) Graphical summary of human CD45^+^ cell proportions of live leukocytes in spleen and ES of mice that were left untreated (untrtd), injected with PBS or free HKCA. n=3-5/group, combined data of 1-3 independent experiments Statistical significance determined by ANOVA and Tuckey’s test for multiple comparison; mean +/-SD. (**c**) The ES of huPBMC-ES-NSG mice were injected with PBS, free HKCA, LPS-activated moDC or HKCA-loaded moDC and harvested 7 days after the last injection similar to (a). Graphical summary of the fraction of Ki67^+^ CD4^+^ cells within the spleen and ES after i.d. injections of the ES as indicated, gated on viable CD45^+^CD3^+^ cells.

**Supplementary Figure 8:**
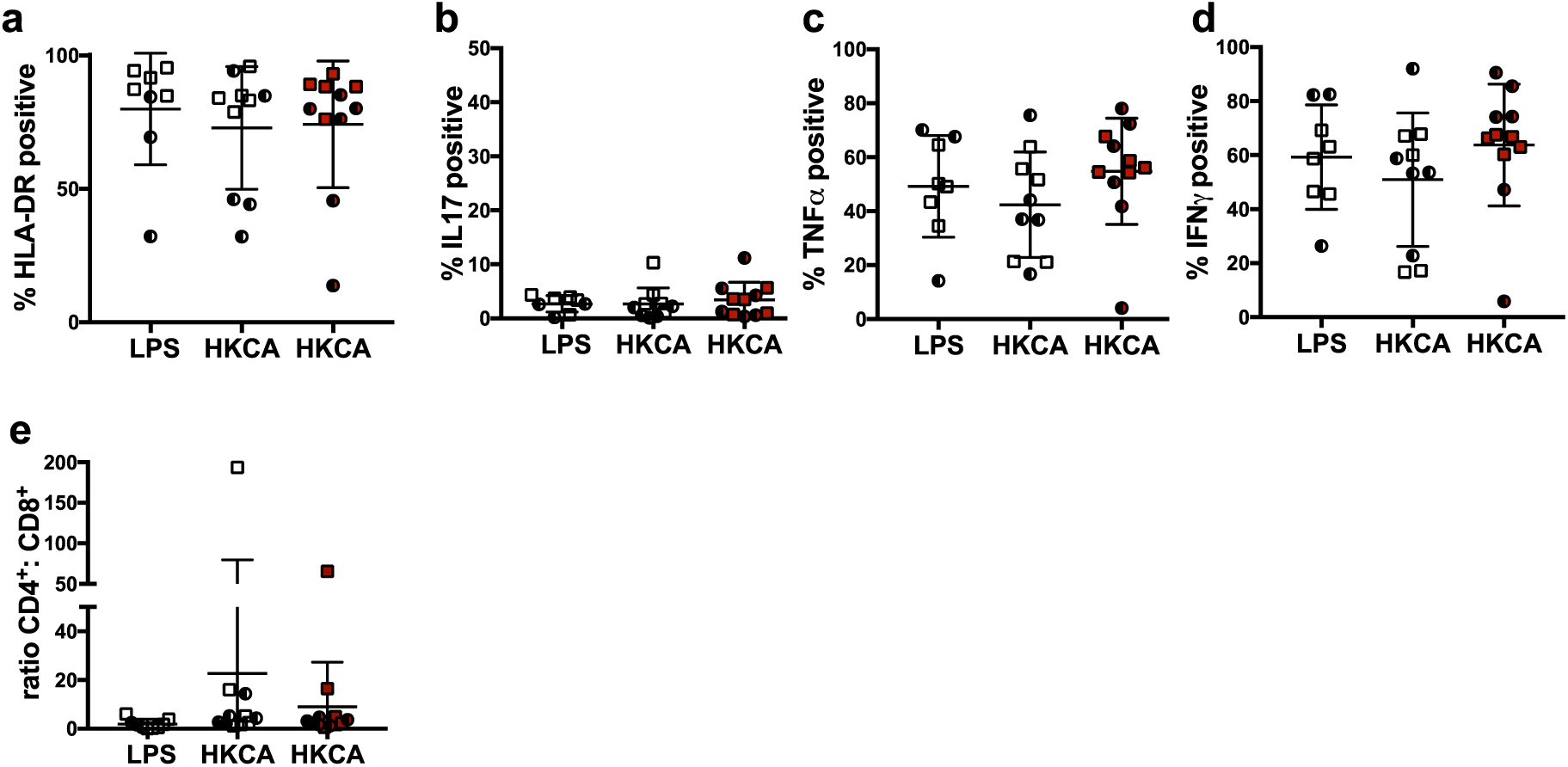
Splenic T cell are not activated by an allogeneic ES or by HKCA presented in the ES. NSG mice bearing fully healed ES of one of two different skin donors (A and B) were adoptively transferred with either skin donor-matched PBMC or skin donor-mismatched PBMC. Intradermal injections of donor A derived LPS/moDC or HKCA/moDC were performed as depicted in Fig. 6 (i.e. leukocytes were matched). Single cell suspensions of spleens were analyzed by flow cytometry after *ex vivo* stimulation with PMA/Ionomycin and intracellular staining. (**a**) Graphical summary of the expression of HLA-DR by CD4^+^ T cells of CD3^+^CD45^+^ live leukocytes isolated from the spleen of huPBMC-ES-NSG mice after intradermal injection of LPS/moDC (LPS) or HKCA/moDC (HKCA) into ES matched (black or empty symbols) or mismatched (red symbols) to the leukocytes. (**a-d**) Splenocytes were stimulated with PMA/ionomycin *ex vivo* and intracellular cytokine production of the indicated cytokines analyzed by flow cytometry. (**e**) Ratio of CD4^+^ to CD8^+^ cells in spleens of huPBMC-ES-NSG mice, gated on CD3^+^CD45^+^ live leukocytes. Each symbol represents a single animal, data compiled from 2-5 independent experiments as indicated by the fillings of the symbols.

**Table S1:** Detailed list of antibodies and reagents.

**Tissue preparation**
| Reagent | Company | Catalog number |
| --- | --- | --- |
| Collagenase Type 4 | Worthington | LS004186 |
| DNAse | Sigma-Aldrich | DN25 |
| RPMI 1640 | Gibco | 31870074 |
| human serum | Sigma-Aldrich | H5667/H4522 |
| Penicillin/streptavidin | Sigma-Aldrich | P0781 |
| L-Glutamine | Gibco | A2916801 |
| NEAA | Gibco | 11140035 |
| Sodium-Pyruvat | Sigma-Aldrich | S8636 |
| β-Mercaptoethanol | Gibco | 31350-010 |
| PBS | Gibco | 14190169 |
| Ficoll Paque Plus | GE-Healthcare | GE17-1440-02 |
| Protease Inhibitor Cocktail | Sigma-Aldrich | P8340 |

**Cellular activation**
| Reagent | Company | Catalog number |
| --- | --- | --- |
| Brefeldin A | Sigma-Aldrich | B6542 |
| Cytofix/Cytoperm kit | BD | RUO 554714 |
| Foxp3 / Transcription Factor Staining Buffer Set | Invitrogen | 00-5523-00 |
| Ionomycin | Sigma-Aldrich | I06434 |
| PMA | Sigma-Aldrich | P8139 |
| Recombinant human IL2 | Immunotools | 11340023 |
| Recombinant human IL4 | Immunotools | 11340047 |
| Recombinant human GM-CSF | Immunotools | 11343127 |
| Purified NA/LE Mouse anti-human CD28 (CD28.2) | BD | 555725 |
| Functional Grade, CD3 monoclonal Antibody (OKT3) | eBioscience | 16-0037-85 |
| Cytokine/Chemokine/Growth Factor 45-Plex Human ProcartaPlex™ | Invitrogen | EPX-450-12171-901 |

**Skin cell culture and transplantation**
| Reagent | Company | Catalog number |
| --- | --- | --- |
| Epilife | Gibco | MEPICF500 |
| DMEM | Gibco | 11960-044 |
| MEM | Gibco | 11380037 |
| TrypLE express | Gibco | 12604021 |
| Primocin | invitrogen | ant-pm-1 |
| X-VIVO 15 | Lonza | 881028 |

**Tissue preparation from mice**
| Reagent | Company | Catalog number |
| --- | --- | --- |
| BD Pharm Lyse | BD | 555899 |
| Collagenase from Clostridium histolyticum | Sigma-Aldrich | C9407 |
| Hyaluronidase | Sigma-Aldrich | H3506 |

**Histology**
| Reagent | Company | Catalog number |
| --- | --- | --- |
| Cell Conditioning 1 (CC1) | Roche | 950-124 |
| Eosin Y aqueous solution | Sigma | HT110232 |
| Hemalum solution acid acc. to Mayer | Carl Rorth | T865.1 |
| Human Cytokeratin 5/6 (D5/16B4) | Roche | 790-4554 |
| ProLong™ Gold Antifade Mountant with DAPI | Invitrogen | P36931 |
| TissueTek O.C.T. Compound | Sakura | TTEK |
| ultraView Universal DAB Detection kit | Roche | 760-500 |

**Antibodies**
| Reagent | Company | Catalog number |
| --- | --- | --- |
| CLA FITC (HECA-452) | Biolegend | 321306 |
| CLA bv605 (HECA-452) | BD | 563960 |
| CD1a FITC (HI149) | BioLegend | 300104 |
| CD3 PerCP-Cy5.5 (OKT3) | eBioscience | 45-0037-42 |
| CD3 BV421 (UCHT1) | BioLegend | 300434 |
| CD3 bv605 (SK7) | BioLegend | 344835 |
| CD3 PE (HIT3a) | BD | 561803 |
| CD3 PE-Cy5.5 (UCHT1) | eBioscience | 15-0038-42 |
| CD4 PE-594 (RPA-T4) | BioLegend | 300548 |
| CD4 PE-Cy5 (RPA-T4) | BioLegend | 300510 |
| CD4 Alexa Fluor 700 (RPA-T4) | BioLegend | 300526 |
| CD4 bv500 (RPA-T4) | BD | 560768 |
| CD8 PE (OKT8) | eBioscience | 12-0086-42 |
| CD8 Pacific Orange (3B5) | eBioscience | MHCD0830 |
| CD8 bv510 (RPA-T8) | BioLegend | 301048 |
| CD14 bv421 (M5E2) | BioLegend | 301830 |
| CD25 PE-Cy5 | BioLegend | 302608 |
| CD25 PE-Cy7 (BC96) | eBioscience | 302612 |
| CD45 PerCP-Cy5.5 (HI30) | BD | 564105 |
| CD45 APC (HI30) | eBioscience | 17-0459-42 |
| CD45 BV785 (HI30) | BioLegend | 304048 |
| CD45RA PE-e610 (HI100) | eBioscience | 61-0458-42 |
| CD45RAPerCP-Cy5.5 (HI100) | BioLegend | 304122 |
| CD45RA Alexa Fluor 700 (HI100) | BioLegend | 304120 |
| CD62L AF700 (DREG-56) | Biolegend | 304820 |
| CD69 PE (FN50) | BioLegend | 310906 |
| CD69 PerCP/eFluor710 (FN50) | eBioscience | 46-0699-42 |
| CD86 PE (B7-2) | eBioscience | 12-0869-41 |
| CCR6 PE-Cy7 (G034E3) | BioLegend | 353418 |
| CCR7 bv605 (G043H7) | BioLegend | 353224 |
| GM-CSF bv421 (BVD2-21C11) | BD | 562930 |
| HLA-DR AF700 (L243) | BioLegend | 307626 |
| HLA-DR eFluor 780 (LN3) | eBioscience | 47-9956-42 |
| IFNg PE-Cy7 (B27) | BioLegend | 506518 |
| IFN-g BV605 (4S.B3) | BD | 563731 |
| IFN-g AF700 (4S.B3) | BioLegend | 502520 |
| IL-2 PE (MQ1-17H12) | eBioscience | 12-7029-42 |
| IL17A APC (BL168) | BioLegend | 512334 |
| IL-17A PerCP-Cy5.5 (eBio64DEC17) | eBioscience | 45-7179-42 |
| IL-13 PE (JES10-5A2) | BD | 554571 |
| IL-22 PE-Cy7 (22URT1) | eBioscience | 25-7229-42 |
| IL-22 PE-PE (22URT1) | eBioscience | 12-7229-42 |
| ki67 bv605 (ki67) | BioLegend | 350521 |
| ki-67 PE-Cy7 (SolA15) | eBioscience | 25-5698-82 |
| TNFa FITC (Mab11) | BioLegend | 502915 |
| TNFa PE-594 (Mab11) | BioLegend | 502946 |
| TCRgd PerCP-eFluor710 (B1.1) | eBioscience | 46-9959 |
| Fixable Viability Dye eFluor™ 780 | eBioscience | 65-0865-14 |
| Fixable Viability Dye eFluor™ 520 | eBioscience | 65-0867-14 |
| eBioscience™ Fixable Viability Dye eFluor™ 506 | eBioscience | 65-0866-14 |
| IF primary antibody: rabbit anti-human NC-1 domain of type VII collagen (LH7.2) | kindly provided by Dr. Alexander Nyström, University of Freiburg, Germany |  |
| IF secondary antibody: goat anti-rabbit A488 | ThermoFisher | A11008 |
| mLy6G (Gr-1) InVivoMab RB6-8C5 | BioXcell | BE0075 |
| Cell Proliferation Dye eFluor™ 450 | ThermoFisher | 65-0842-85 |

## References

1. Bos, J. D. et al. Predominance of ‘memory’ T cells (CD4+, CDw29+) over ‘naive’ T cells (CD4+, CD45R+) in both normal and diseased human skin. Arch. Dermatol. Res. 281, 24–30 (1989).

2. Clark, R. A. et al. The vast majority of CLA+ T cells are resident in normal skin. J. Immunol. Baltim. Md 1950 176, 4431–4439 (2006).

3. Nestle, F. O., Di Meglio, P., Qin, J.-Z. & Nickoloff, B. J. Skin immune sentinels in health and disease. Nat. Rev. Immunol. 9, 679–691 (2009).

4. Di Meglio, P., Perera, G. K. & Nestle, F. O. The multitasking organ: recent insights into skin immune function. Immunity 35, 857–869 (2011).

5. Klicznik, M. M., Szenes-Nagy, A. B., Campbell, D. J. & Gratz, I. K. Taking the lead - how keratinocytes orchestrate skin T cell immunity. Immunol. Lett. 200, 43–51 (2018).

6. Gebhardt, T., Palendira, U., Tscharke, D. C. & Bedoui, S. Tissue-resident memory T cells in tissue homeostasis, persistent infection, and cancer surveillance. Immunol. Rev. 283, 54–76 (2018).

7. Masopust, D. & Soerens, A. G. Tissue-Resident T Cells and Other Resident Leukocytes. Annu. Rev. Immunol. 37, null (2019).

8. Snyder, M. E. et al. Generation and persistence of human tissue-resident memory T cells in lung transplantation. Sci. Immunol. 4, eaav5581 (2019).

9. Park, S. L. et al. Tissue-resident memory CD8+ T cells promote melanoma-immune equilibrium in skin. Nature 565, 366–371 (2019).

10. Campbell, J. J., Clark, R. A., Watanabe, R. & Kupper, T. S. Sézary syndrome and mycosis fungoides arise from distinct T-cell subsets: a biologic rationale for their distinct clinical behaviors. Blood 116, 767–771 (2010).

11. Clark, R. A. et al. Skin effector memory T cells do not recirculate and provide immune protection in alemtuzumab-treated CTCL patients. Sci. Transl. Med. 4, 117ra7 (2012).

12. Casey, K. A. et al. Antigen independent differentiation and maintenance of effector-like resident memory T cells in tissues. J. Immunol. Baltim. Md 1950 188, 4866–4875 (2012).

13. Gebhardt, T. et al. Memory T cells in nonlymphoid tissue that provide enhanced local immunity during infection with herpes simplex virus. Nat. Immunol. 10, 524–530 (2009).

14. Glennie, N. D. et al. Skin-resident memory CD4+ T cells enhance protection against Leishmania major infection. J. Exp. Med. 212, 1405–1414 (2015).

15. Mackay, L. K. et al. The developmental pathway for CD103(+)CD8+ tissue-resident memory T cells of skin. Nat. Immunol. 14, 1294–1301 (2013).

16. Park, C. O. et al. Staged development of long-lived T-cell receptor αβ TH17 resident memory T-cell population to Candida albicans after skin infection. J. Allergy Clin. Immunol. 142, 647–662 (2018).

17. Mackay, L. K. et al. T-box Transcription Factors Combine with the Cytokines TGF-β and IL-15 to Control Tissue-Resident Memory T Cell Fate. Immunity 43, 1101–1111 (2015).

18. Pan, Y. et al. Survival of tissue-resident memory T cells requires exogenous lipid uptake and metabolism. Nature 543, 252–256 (2017).

19. Hu, W. & Pasare, C. Location, location, location: tissue-specific regulation of immune responses. J. Leukoc. Biol. 94, 409–421 (2013).

20. McCully, M. L. et al. Epidermis instructs skin homing receptor expression in human T cells. Blood 120, 4591–4598 (2012).

21. Kumar, B. V. et al. Human Tissue-Resident Memory T Cells Are Defined by Core Transcriptional and Functional Signatures in Lymphoid and Mucosal Sites. Cell Rep. 20, 2921–2934 (2017).

22. Gudjonsson, J. E., Johnston, A., Dyson, M., Valdimarsson, H. & Elder, J. T. Mouse models of psoriasis. J. Invest. Dermatol. 127, 1292–1308 (2007).

23. Shay, T. et al. Conservation and divergence in the transcriptional programs of the human and mouse immune systems. Proc. Natl. Acad. Sci. 110, 2946–2951 (2013).

24. Perlman, R. L. Mouse models of human disease. Evol. Med. Public Health 2016, 170–176 (2016).

25. Watanabe, R. et al. Human skin is protected by four functionally and phenotypically discrete populations of resident and recirculating memory T cells. Sci. Transl. Med. 7, 279ra39 (2015).

26. King, M. A. et al. Human peripheral blood leucocyte non-obese diabetic-severe combined immunodeficiency interleukin-2 receptor gamma chain gene mouse model of xenogeneic graft-versus-host-like disease and the role of host major histocompatibility complex. Clin. Exp. Immunol. 157, 104–118 (2009).

27. Boyman, O. et al. Spontaneous Development of Psoriasis in a New Animal Model Shows an Essential Role for Resident T Cells and Tumor Necrosis Factor-α. J. Exp. Med. 199, 731–736 (2004).

28. Racki, W. J. et al. NOD-scid IL2rgamma(null) mouse model of human skin transplantation and allograft rejection. Transplantation 89, 527–536 (2010).

29. Sanchez Rodriguez, R. et al. Memory regulatory T cells reside in human skin. J. Clin. Invest. 124, 1027–1036 (2014).

30. Carretero, M. et al. Differential Features between Chronic Skin Inflammatory Diseases Revealed in Skin-Humanized Psoriasis and Atopic Dermatitis Mouse Models. J. Invest. Dermatol. 136, 136–145 (2016).

31. Guerrero-Aspizua, S. et al. Development of a Bioengineered Skin-Humanized Mouse Model for Psoriasis. Am. J. Pathol. 177, 3112–3124 (2010).

32. Boyce, S. T. et al. Skin anatomy and antigen expression after burn wound closure with composite grafts of cultured skin cells and biopolymers. Plast. Reconstr. Surg. 91, 632–641 (1993).

33. Burke, J. F., Yannas, I. V., Quinby, W. C., Bondoc, C. C. & Jung, W. K. Successful use of a physiologically acceptable artificial skin in the treatment of extensive burn injury. Ann. Surg. 194, 413–428 (1981).

34. Nanchahal, J. & Davies, D. Cultured composite skin grafts for burns. BMJ 301, 1342–1343 (1990).

35. Clark, R. A. Resident memory T cells in human health and disease. Sci. Transl. Med. 7, 269rv1 (2015).

36. Jiang, X. et al. Skin infection generates non-migratory memory CD8+ T(RM) cells providing global skin immunity. Nature 483, 227–231 (2012).

37. Campbell, J. J. & Butcher, E. C. Chemokines in tissue-specific and microenvironment-specific lymphocyte homing. Curr. Opin. Immunol. 12, 336–341 (2000).

38. Nowak, K. et al. Absence of γ-Chain in Keratinocytes Alters Chemokine Secretion, Resulting in Reduced Immune Cell Recruitment. J. Invest. Dermatol. 137, 2120–2130 (2017).

39. Uchi, H., Terao, H., Koga, T. & Furue, M. Cytokines and chemokines in the epidermis. J. Dermatol. Sci. 24, S29–S38 (2000).

40. Merkley, M. A. et al. Large-scale analysis of protein expression changes in human keratinocytes immortalized by human papilloma virus type 16 E6 and E7 oncogenes. Proteome Sci. 7, 29 (2009).

41. Wang, C. K., Nelson, C. F., Brinkman, A. M., Miller, A. C. & Hoeffler, W. K. Spontaneous cell sorting of fibroblasts and keratinocytes creates an organotypic human skin equivalent. J. Invest. Dermatol. 114, 674–680 (2000).

42. Wetzels, R. H. W., Robben, H. C. M., Leigh, I. M., Vooijs, G. P. & Ramaekerst, F. C. S. Distribution Patterns of Type VII Collagen in Normal and Malignant Human Tissues. 139, 9 (1991).

43. Ali, N. et al. Xenogeneic Graft-versus-Host-Disease in NOD-scid IL-2Rγnull Mice Display a T-Effector Memory Phenotype. PLoS ONE 7, (2012).

44. King, M. et al. A new Hu-PBL model for the study of human islet alloreactivity based on NOD-scid mice bearing a targeted mutation in the IL-2 receptor gamma chain gene. Clin. Immunol. Orlando Fla 126, 303–314 (2008).

45. Shultz, L. D. et al. Human Lymphoid and Myeloid Cell Development in NOD/LtSz-scid IL2Rγnull Mice Engrafted with Mobilized Human Hemopoietic Stem Cells. J. Immunol. 174, 6477–6489 (2005).

46. Carr, M. W., Roth, S. J., Luther, E., Rose, S. S. & Springer, T. A. Monocyte chemoattractant protein 1 acts as a T-lymphocyte chemoattractant. Proc. Natl. Acad. Sci. U. S. A. 91, 3652–3656 (1994).

47. Kawai, T. et al. Selective diapedesis of Th1 cells induced by endothelial cell RANTES. J. Immunol. Baltim. Md 1950 163, 3269–3278 (1999).

48. Fukui, A. et al. Interleukin-8 and CXCL10 expression in oral keratinocytes and fibroblasts via Toll-like receptors. Microbiol. Immunol. 57, 198–206 (2013).

49. Nanki, T. & Lipsky, P. E. Stimulation of T-Cell Activation by CXCL12/Stromal Cell Derived Factor-1 Involves a G-Protein Mediated Signaling Pathway. Cell. Immunol. 214, 145–154 (2001).

50. Moser, B. & McCully, M. L. The Human Cutaneous Chemokine System. Front. Immunol. 2, (2011).

51. Adachi, T. et al. Hair follicle-derived IL-7 and IL-15 mediate skin-resident memory T cell homeostasis and lymphoma. Nat. Med. 21, 1272–1279 (2015).

52. Belarif, L. et al. IL-7 receptor blockade blunts antigen-specific memory T cell responses and chronic inflammation in primates. Nat. Commun. 9, (2018).

53. Wang, X. et al. Engraftment of human central memory-derived effector CD8+ T cells in immunodeficient mice. Blood 117, 1888–1898 (2011).

54. Klicznik, M. M. et al. Human CD4+CD103+ cutaneous resident memory T cells are found in the circulation of healthy individuals. Sci. Immunol. 4, (2019).

55. Shannon, M. F., Himes, S. R. & Coles, L. S. GM-CSF and IL-2 share common control mechanisms in response to costimulatory signals in T cells. J. Leukoc. Biol. 57, 767–773 (1995).

56. Shi, Y. et al. Granulocyte-macrophage colony-stimulating factor (GM-CSF) and T-cell responses: what we do and don’t know. Cell Res. 16, 126–133 (2006).

57. Belkaid, Y. & Tamoutounour, S. The influence of skin microorganisms on cutaneous immunity. Nat. Rev. Immunol. 16, 353–366 (2016).

58. Gosselin, A. et al. Peripheral blood CCR4+CCR6+ and CXCR3+CCR6+CD4+ T cells are highly permissive to HIV-1 infection. J. Immunol. Baltim. Md 1950 184, 1604–1616 (2010).

59. Klein, R. S. et al. Oral candidiasis in high-risk patients as the initial manifestation of the acquired immunodeficiency syndrome. N. Engl. J. Med. 311, 354–358 (1984).

60. Lagunes, L. & Rello, J. Invasive candidiasis: from mycobiome to infection, therapy, and prevention. Eur. J. Clin. Microbiol. Infect. Dis. Off. Publ. Eur. Soc. Clin. Microbiol. 35, 1221–1226 (2016).

61. Ling, Y. et al. Inherited IL-17RC deficiency in patients with chronic mucocutaneous candidiasis. J. Exp. Med. 212, 619–631 (2015).

62. Puel, A. et al. Chronic mucocutaneous candidiasis in humans with inborn errors of interleukin-17 immunity. Science 332, 65–68 (2011).

63. Acosta-Rodriguez, E. V. et al. Surface phenotype and antigenic specificity of human interleukin 17-producing T helper memory cells. Nat. Immunol. 8, 639–646 (2007).

64. Hernández-Santos, N. & Gaffen, S. L. Th17 cells in immunity to Candida albicans. Cell Host Microbe 11, 425–435 (2012).

65. Holling, T. M., Schooten, E. & van Den Elsen, P. J. Function and regulation of MHC class II molecules in T-lymphocytes: of mice and men. Hum. Immunol. 65, 282–290 (2004).

66. Oshima, S. & Eckels, D. D. Selective signal transduction through the CD3 or CD2 complex is required for class II MHC expression by human T cells. J. Immunol. Baltim. Md 1950 145, 4018–4025 (1990).

67. Ko, H. S. Ia determinants on stimulated human T lymphocytes. Occurrence on mitogen- and antigen-activated T cells. J. Exp. Med. 150, 246–255 (1979).

68. Duhen, T., Geiger, R., Jarrossay, D., Lanzavecchia, A. & Sallusto, F. Production of interleukin 22 but not interleukin 17 by a subset of human skin-homing memory T cells. Nat. Immunol. 10, 857–863 (2009).

69. Schlapbach, C. et al. Human TH9 cells are skin-tropic and have autocrine and paracrine proinflammatory capacity. Sci. Transl. Med. 6, 219ra8–219ra8 (2014).

70. Zielinski, C. E. et al. Pathogen-induced human TH17 cells produce IFN-γ or IL-10 and are regulated by IL-1β. Nature 484, 514–518 (2012).

71. Kawai, T. et al. Selective diapedesis of Th1 cells induced by endothelial cell RANTES. J. Immunol. Baltim. Md 1950 163, 3269–3278 (1999).

72. Fidan, I. et al. The effects of fluconazole and cytokines on human mononuclear cells. Mem. Inst. Oswaldo Cruz 102, 127–131 (2007).

73. Zuber, J. et al. Bidirectional intragraft alloreactivity drives the repopulation of human intestinal allografts and correlates with clinical outcome. Sci. Immunol. 1, (2016).

74. Fenini, G. et al. Genome Editing of Human Primary Keratinocytes by CRISPR/Cas9 Reveals an Essential Role of the NLRP1 Inflammasome in UVB Sensing. J. Invest. Dermatol. 138, 2644–2652 (2018).

75. Farber, D. L., Yudanin, N. A. & Restifo, N. P. Human memory T cells: generation, compartmentalization and homeostasis. Nat. Rev. Immunol. 14, 24–35 (2014).

76. Saule, P. et al. Accumulation of memory T cells from childhood to old age: central and effector memory cells in CD4(+) versus effector memory and terminally differentiated memory cells in CD8(+) compartment. Mech. Ageing Dev. 127, 274–281 (2006).

77. Haynes, L. & Swain, S. L. Why Aging T Cells Fail: Implications for Vaccination. Immunity 24, 663–666 (2006).

78. Gratz, I. K. et al. Cutting edge: Self-antigen controls the balance between effector and regulatory T cells in peripheral tissues. J. Immunol. Baltim. Md 1950 192, 1351–1355 (2014).

79. Sallusto, F. & Lanzavecchia, A. Efficient presentation of soluble antigen by cultured human dendritic cells is maintained by granulocyte/macrophage colony-stimulating factor plus interleukin 4 and downregulated by tumor necrosis factor alpha. J. Exp. Med. 179, 1109–1118 (1994).

